# Dual-topology of collagen XV and tenascin C acts in concert to guide and shape developing motor axons

**DOI:** 10.1101/2023.05.18.541309

**Authors:** Laurie Nemoz-Billet, Martial Balland, Laurent Gilquin, Benjamin Gillet, Isabelle Stévant, Emilie Guillon, Sandrine Hughes, Gilles Carpentier, Elisabeth Vaganay, Mary-Julieth Gonzalez-Melo, Manuel Koch, Yad Ghavi-Helm, Florence Ruggiero, Sandrine Bretaud

## Abstract

During development, motor axons are guided towards their muscle target by various extrinsic cues including extracellular matrix (ECM) proteins those identities remain poorly documented. Using single-cell RNA-sequencing of differentiating slow muscle progenitors (SMP) in zebrafish, we charaterized the SMP as a major source of ECM proteins that were computationally predicted to form a basement membrane-like structure tailored for motor axon guidance. Multiple *in vivo* and *in vitro* approaches further revealed that motor axon shape and growth relies on the timely expression of the attractive cue Collagen XV-B (ColXV-B) that locally provides motor axons with a permissive soft microenvironment and separately organizes the repulsive cue Tenascin C into a unique functional dual topology. Bioprinted micropatterns mimicking their unique topology provide compelling evidence that it represents a sufficient condition to elicit directional motor axon growth. Our study provides the first evidence that ECM topology and stiffness critically influence motor axon navigation in vertebrates with potential applications in regenerative medicine for peripheral nerve injury.

## Introduction

During development, axons of motor neurons follow stereotypical trajectories to their muscle targets, guided by various molecular cues. The extracellular matrix (ECM) can influence motor axon navigation by providing external chemical and mechanical guidance cues to developing axons, attracting or repelling them. Among them are the well-known diffusible chemical cues, netrins and semaphorins, that likely collaborate with other extracellular cues to guide developing axons (Dickson et al, 2002). Yet, contact guidance cues, which are immobilized ECM proteins, remain poorly documented. The zebrafish has been proven to be a powerful animal model to investigate peripheral nervous system development and more specifically motor axon navigation (Beattie, 2000). There are 3 types of primary motor neurons per spinal cord hemisegment that are identified by their cell body position in the spinal cord: the rostral primary (RoP), the middle primary (MiP) and the caudal primary (CaP). At 17 hours post fertilization (hpf), primary motor axons initiate outgrowth and exit the spinal cord, a process that is pioneered by the CaP. They next project through a common path and pause at the choice point which corresponds to an intermediate target located at the level of the horizontal myosepta. Then, the RoP, MiP and Cap axons diverge to innervate specific territories, respectively the lateral, dorsal and ventral myotomes. The myotome is an important source of guidance cues for developing motor axons (Beattie, 2000). In particular, because of their position in the trajectory of the future growing axons, the slow muscle precursors (SMPs) (a.k.a adaxial cells) and muscle pioneers (MPs) have been shown to play a pivotal role in motor axon navigation (Melançon et al 1997, Zhang and Granato, 2000). As the myotome develops and before motor axon extension, SMPs stack, elongate and traverse the myotome to reach its periphery where they eventually differentiate into slow superficial fibers (Daggett et al, 2007). Only a subset of SMPs differentiate into MPs and remain located adjacent to the notochord (Stickney et al, 2000). As motor axon exit from the spinal cord, they extend through an ECM path mainly produced and deposited locally by SMPs that were identified as the major source of ECM guidance cues (Beattie, 2000).

ECM forms complex and intricate protein networks whose composition and spatial organisation are closely related to their function in tissues (Ricard-Blum and Ruggiero, 2005). Chondroitin sulfate proteoglycans and Tenascin-C (TnC) are both expressed by SMP and have been identified as important guidance cues for zebrafish motor axons (Bernhardt and Schachner 2000, Zhang et al 2004, Schweitzer et al 2005). Notably, TnC morphants display abnormal extra-branching axons indicative of a role of TnC as a repulsive local molecular cue that guides away motor axons (Schweitzer et al 2005). Collagens also play a role in axon navigation in the zebrafish central nervous system, as well as in the peripheral nervous system (reviewed in Bretaud et al, 2019). Defects in motor axon navigation were observed both in Collagen XVIII morphants (Schneider and Granato, 2006) and Collagen XIX mutants (*stumpy* mutants) (Hilario et al 2010). More recently, we identified Collagen XV-B (ColXV-B) as an attractive contact guidance cue. We found that ColXV-B is deposited in the motor path by the SMPs before they migrate at the myotome periphery through a unique two-step mechanism involving the notochord Shh and myotomal MuSK signaling (Guillon et al, 2016). In invertebrates, the absence of the ColXV/ColXVIII orthologs, Dmp/Mp in drosophila and cle-1 in *C. elegans*, also led to defects in motor axon navigation (Meyer and Moussian, 2009; Ackley et al, 2001), suggesting that the role of ColXV in axonogenesis is well conserved across species. The Collagen XV and ColXVIII that are structurally closely related and both belong to the subgroup of multiplexins, as well as the FACIT Collagen XIX are all described as basement membrane zone (BMZ)-associated collagens suggesting that the motor axon trajectory consists in a highly specialized BMZ. Yet, although a few ECM molecules have been described as contact guidance cues, the full composition and specifically their organization in the ECM motor path remain surprisingly unresolved. To fill this gap, we first performed a single cell RNA sequencing (scRNAseq) analysis of differentiating SMPs to characterize the matrisome of SMPs. Bioinformatics analysis of the data show that differentiated SMPs both express BM-toolkit genes and specific core-matrisome genes involved in motor axon navigation that together can organize into a BMZ-like structure tailored for motor axon guidance. In support of this, image analysis of immunofluorescence staining combined with loss of function experiments and biomechanics and bioengineering analysis unveil that ColXV-B, the earliest core-matrisome gene expressed by differentiated SMPs, creates a favorable microenvironment for axon growth and navigation by influencing ECM motor path stiffness and topology. Our study put forward the paracrine role of SMP in shaping ECM path critical for future motor axon navigation towards their muscle target.

## Results

### Single-cell profiling unravels the dynamics of SMP differentiation

Before their final differentiation into slow superficial fibres, SMPs undergo a series of morphological changes, from pseudo-epithelial to elongated pre-migratory, as the myotome matures (Fig. 1A). Strikingly, while the SMP stereotypical behaviours are well characterized, their related-gene signatures remain unknown. We thus first aimed at characterizing the trajectory and gene signatures of SMP differentiation using scRNAseq. With this aim, we took advantage of the rostro-caudal polarity of the segmented somites and the segmentation clock during which one additional somite is formed every 30 mins to isolate SMPs at different stages of their differentiation in the same embryo. One-cell stage embryos were injected with *smyhc1:gfp* construct and the cell suspension isolated from 18 hpf trunk injected embryos was used to sort GFP-positive cells and perform scRNAseq (Fig. 1B). Unbiased clustering analysis of our scRNAseq data revealed 18 clusters. Four clusters were enriched for cells that expressed *smyhc1* and *prdm1a*, two specific markers of the slow muscle lineage: cluster 1 (491 cells), cluster 5 (347 cells), cluster 10 (175 cells) and cluster 11 (172 cells) (Fig. 1C). The mapping against the zebrafish genome containing the *gfp* gene indicates that *smyhc1* expressing cells exhibit a high expression of *gfp* which confirms the SMP identity of *smyhc1* positive cells (Supplementary Fig1). We thus further focused on the analysis of these four clusters annotated as slow muscle lineage derived-cells.

**Figure 1.**
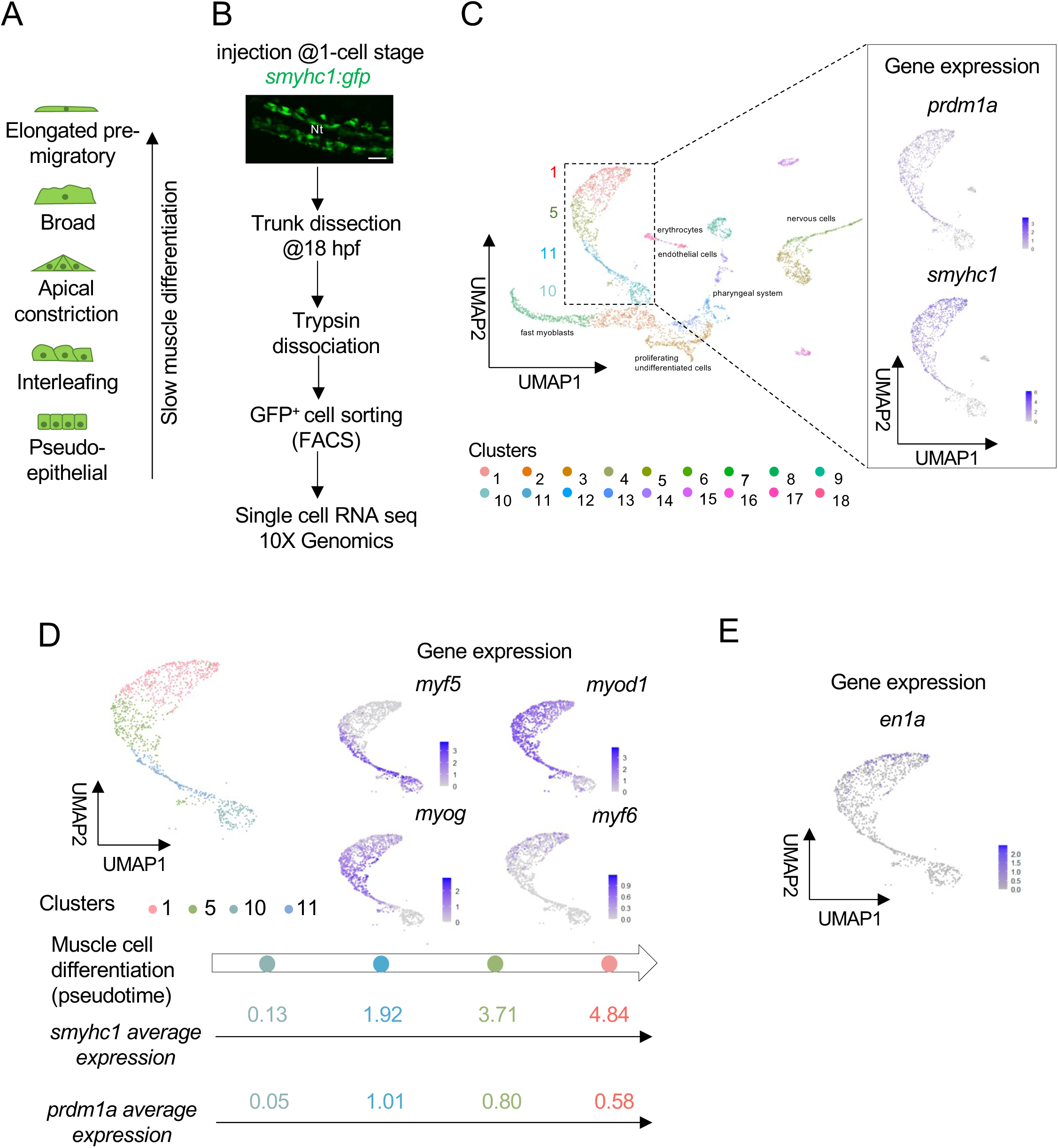
Differentiating SMPs display specific gene expression signatures. (**A**) Schematics of the morphological series of differentiating SMPs in 18hpf embryos; **(B)** Schematic representation of the protocol used for scRNAseq analysis; Confocal projection of differentiating SMP revealed by mosaic *smyhc1:gfp* expression in a living 18 hpf embryo. Dorsal view. Scale bar = 50 µm. N, notochord. Anterior is left. **(C)** UMAP plot obtained after mapping on the zebrafish genome and clusterisation of isolated GFP^+^ cells (left panel); UMAP feature plots showing expression patterns of slow muscle lineage markers *prdm1a* and *smyhc1* (right panel). **(D)** UMAP plot obtained after isolation of the clusters corresponding to SMP lineage; UMAP feature plots showing the expression pattern of myogenic regulatory factors in differentiating SMPs (right panel); Pseudotime trajectory of differentiating SMP and *smyhc1* and *prdm1a* average expression in each cluster (bottom panel). Colors referred to as different clusters. (E) UMAP feature plot showing the expression of the MP marker *en1a* in differentiating SMPs.

We next decided to construct the pseudo-temporal SMP differentiation trajectory based on the expression of myogenic regulatory factors that are known to be differentially expressed during SMP differentiation. The increasing level of *smyhc1* in each of the 4 clusters confirmed their level of maturation (Fig. 1D). Previous *in situ* analyses showed that the expression of *prdm1a* in tail-bud stage and 18 hpf embryos begins at the interleafing / pseudo-epithelial SMP stages. It then reaches a peak of expression in apical constriction / broad SMP stages and decreases in pre-migrating myoblasts of anterior somites (Baxendale et al, 2004). Together, these temporal and spatial myogenic gene expression data suggest that the less mature myoblasts present in cluster 10 might correspond to interleafing / pseudo-epithelial SMP stages while the pre-migrating cells are present in cluster 1. Continuing this deductive way of reasoning, we conclude that the peak of *prdm1a* expression observed in cluster 11 cells corresponds to the apical constriction / broad SMP stages and that the decrease in *prdm1a* expression observed in cluster 5 intermediate SMPs corresponds to the elongated pre-migratory stage (Fig. 1D). As a confirmation of the robustness of the method used to allocate SMPs trajectory to the different clusters, our scRNAseq data clearly subclusterized a subpopulation of 70 cells in the pre-migrating cluster that are identified as muscle pioneers (MP) cells as they additionally and specifically expressed the MP *engrailed1a* (*en1a*) marker (Fig. 1E). In conclusion, cluster 10 corresponds to the less differentiated myoblasts and cluster 1 to the most differentiated ones while clusters 11 and 5 respectively comprise cells of successive intermediate stages of differentiation. By resolving SMPs pseudotemporal trajectory at single cell level, our data uniquely correlate the differentiating SMPs morphological signatures to specific gene expression signatures.

### Differentiating SMP express BM-like specialized ECM genes

We further aimed at characterizing the ECM gene expression landscape of differentiating SMPs. Using our in-silico zebrafish matrisome database (Nauroy et al, 2018), we determined that SMPs globally express 9% of total zebrafish matrisome genes (91/1004). Among these ECM genes and independently of the differentiating stages, SMPs expressed both core matrisome genes (36/335) and matrisome-associated genes (55/669) which comprise ECM-affiliated genes, ECM regulators and secreted factors (Fig. 2A, Supplementary Table 1-4). The less differentiated SMPs (cluster 10) mainly express matrisome-associated genes (34 genes among 51 matrisome genes), especially ECM regulators and secreted factors (13 and 17, respectively). Notably, expression of ECM regulators such as *mmp2* and *mmp14a* in cluster 10 cells indicate an ECM remodelling of the cell microenvironment. On the contrary, elongated pre-migratory SMPs (cluster 5) predominantly express core matrisome genes (29 genes among 56 matrisome genes), suggesting that these SMPs are the main cell source of ECM cues (Fig. 2A). Based on the GO term “axon guidance” (GO:0007411) and data from the literature, we found that SMPs, independently of their differentiating stages, express core matrisome genes and matrisome-associated genes that are known to be involved in axon guidance such as *agrn*, *col15a1b*, *col18a1a*, *col19a1*, *lama5*, *ntn1a*, and *tnc*; and the classic guidance cues *cxcl12a*, *cxcl12b*, *plxna3*, *plxnb1a*, *sema4ba* and *vegfaa* (Fig. 2A, Supplementary Table 1-4). BM remodeling is also shown to guide neurons mainly by locally immobilizing diffusible molecular cues such as netrin and semaphorin (reviewed in Sherwood, 2021). In this way, some of the classic guidance cues that were found to be expressed by SMP could be involved in motor axon pathfinding through their binding to BM proteins.

**Figure 2.**
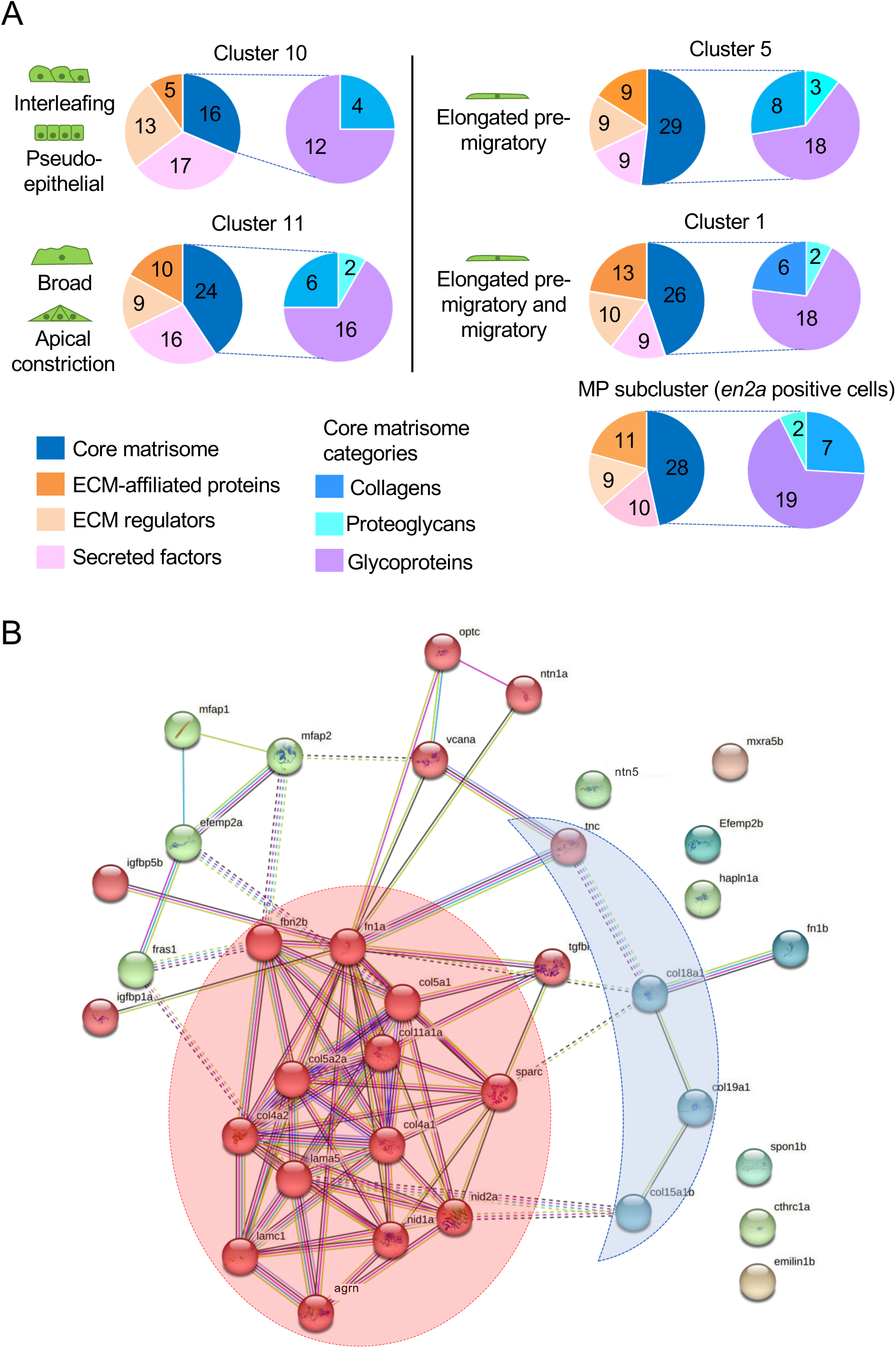
Characterization of differentiating SMP matrisomes. (**A**) Left pie charts represent the number of matrisome genes expressed in each cluster and classified by categories (core-matrisome, ECM-affiliated proteins, ECM regulators, secreted factors), right pie charts represent the number of core-matrisome genes expressed in each cluster and by sub-categories (collagens, proteoglycans, glycoproteins). **(B)** Protein-protein interaction network analysis of the total core matrisome genes expressed by SMPs with STRING database. The protein interaction network of the 36 core-matrisome genes was created with an enrichment p-value <10^-16^. Clusters were generated using a MCL inflation parameter of 2. One color represents one cluster. 2 hubs of interest are highlighted: BM-toolkit (red) and BM-associated genes (blue).

Finally, we characterized the matrisome of the MP cells that are described as intermediate targets for growing axons (Melançon et al, 1997). As the more mature SMPs (clusters 1 and 5), MP predominantly express core matrisome genes, that only differ from SMPs core matrisome genes by 4 genes: *col1a2, colq, lgi1a and vwa2* (Supplementary Table 5). STRING based protein interactions was used to build the interactome of the total core matrisome genes expressed by SMPs. This interaction network reveals a major hub (in red) composed of 20 genes including those encoding the toolkit BM proteins collagen IV, laminins and nidogens, suggesting that the motor axon path ECM mainly consists in a specialized BM. Interestingly, STRING analysis and clustering also unravel a crescent-shaped protein interactions between the main hub and BM-associated proteins all previously described as ECM guidance cues, TnC, ColXVIII, ColXIX and ColXV-B (Hilario et al, 2010, Schweitzer et al, 2005, Schneider and Granato 2006, Guillon et al, 2016) (Fig. 2B). Our data show that prior their migration to reach the final destination at the periphery of the myotome, SMPs timely express genes encoding BM genes and BM-associated axon guidance local cues. These ECM proteins are likely to assemble into a unique BM-like structure and organisation that is required for its specialized function.

### The TnC and ColXV-B guidance cues exhibit dual topology that confines the motor axon trajectory

To test this hypothesis, we decided to visualize the spatial organisation of the BM-associated proteins with confocal microscopy. *tnc*, *col18a1* and *col15a1b* all start to be expressed in cluster 11 (Supplementary Table 2) while the onset of *col19a1* expression occurs in cluster 5 (Supplementary Table 3). As TnC and ColXV-B exhibit functional duality (Schweitzer et al, 2005; Guillon et al, 2016), we reasoned that these two immobilized guidance cues organize in a specific way to appropriately guide motor axons. We thus performed double immunostaining of 27 hpf *mnx1:gfp* embryos to directly visualize trunk motor axons with our home-made guinea-pig antibodies to ColXV-B (Supplementary Fig 2) and rabbit antibodies to Tenascin C. Stunningly, our double immunostaining unveils that TnC and ColXV-B display distinctive and even exclusive spatial expression patterns. While ColXV-B is deposited along the motor axon path in a ladder fashion (Fig. 3A) as we previously showed (Guillon et al, 2016), TnC is restricted to the choice point where it forms a channel that delineated the motor axon path (Fig. 3A). Orthogonal views confirm the dual topology of the ColXV-B/TnC deposits (Fig. 3A). Whereas ColXV-B intermittently wraps motor axon all along its trajectory, TnC forms borders which vertically delimit the motor axon path (Fig. 3A). We additionally show that both proteins are regulated by Shh signaling that is required for commitment of SMPs (Supplementary Fig. 3; Bretaud et al, 2011; Guillon et al, 2016). We thus conclude that SMPs timely produce the duo of repulsive and attractive ECM guidance cues, TnC and ColXV-B that are spatially organized to control locally motor axon trajectory.

**Figure 3.**
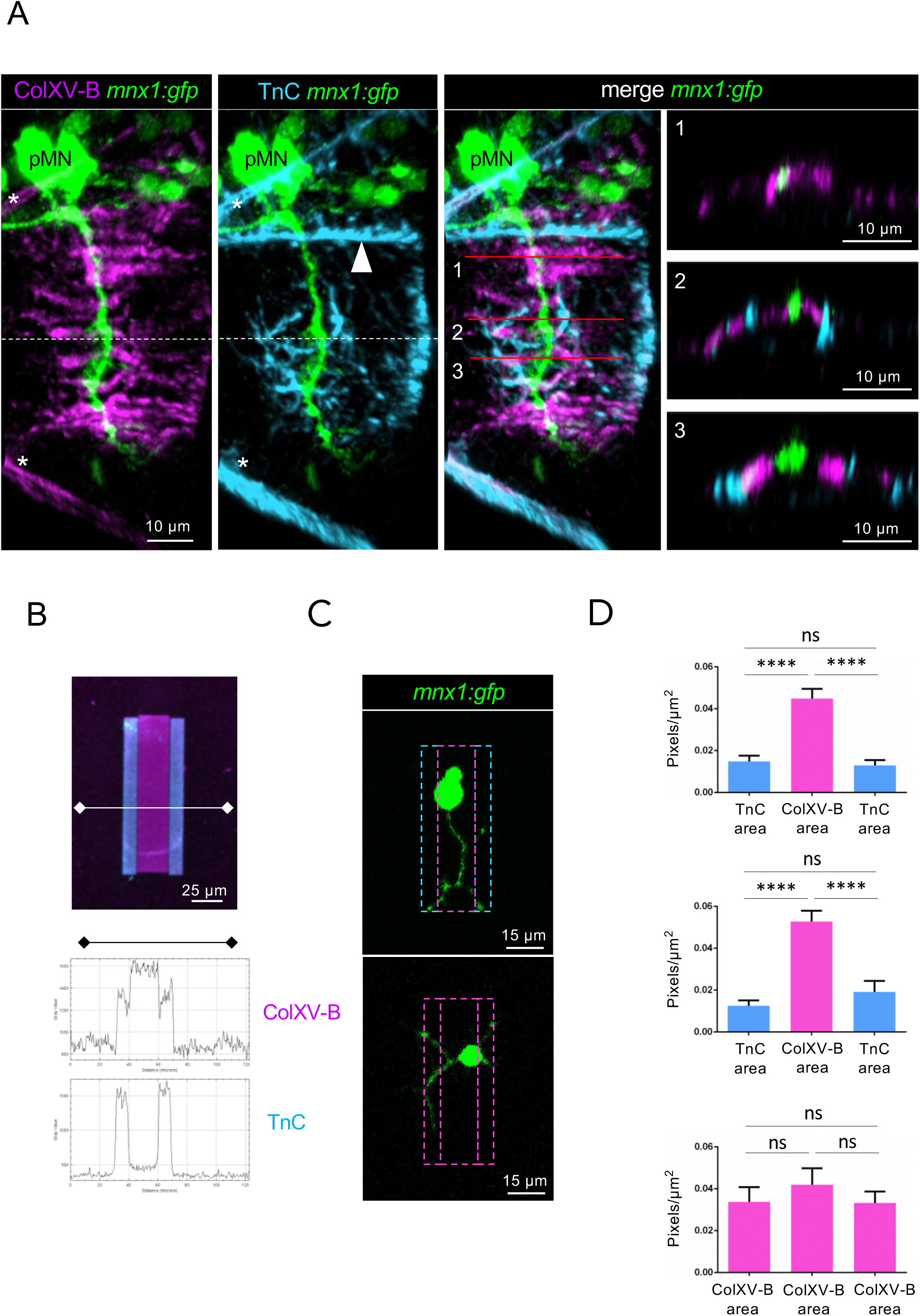
The motor axon trajectory is formed by a central ColXV-B path delimited by TnC borders. (**A**) 3D Imaris images of whole-mount immunostaining of 27 hpf *mnx1:gfp* embryos with anti-rNC1 (magenta, ColXV-B), anti-TnC (cyan), and anti-GFP antibodies (green, motor neurons). Left panel, lateral views; right panel, orthogonal views; anterior is left. Dotted line shows the horizontal myosepta. Asterisks indicate the vertical myosepta. Arrowhead points to the dorsal edge of the notochord. Numbers indicate the level of virtual orthogonal sections **(B)** Double immunostaining of 45×70 µm ColXV-B / TnC double micropatterns with anti-rNC1 (magenta, ColXV-B) and anti-TnC (cyan), plot profile of ColXV-B and TnC staining (Fiji software). **(C)** Live confocal images of motor neurons and axons 24 h after seeding on 31×70 µm ColXV-B / TnC double micropatterns (top panel) and ColXV-B control micropatterns (bottom panel). **(D)** Quantification of the number of pixels corresponding to the neurites for each ColXV-B or TnC area expressed in percentage of the total micropattern area. Top panel, ColXV-B / recombinant murine TnC double micropatterns; n=30; Middle panel, ColXV-B / human purified TnC double micropatterns, n=21; Bottom panel, ColXV-B control micropatterns, n=23. Statistical analysis was performed using a one-way ANOVA. ****p<0.0001; ns: non-significant. Error bars are SEM.

To verify if this topology is decisive for motor axon shaping and navigation, we reproduce the dual topology of TnC and ColXV-B using the Alveole lab technology. With this aim, we used recombinant ColXV-B and TnC proteins produced in HEK293 cells (Supplementary Fig. 2A). The proper organization of the resulting double bioprinted ECM micropatterns was assessed by immunostaining and unveils the expected topology: ColXV-B in the central route delineated by TnC deposits (Fig. 3B). Micropatterns in which TnC borders were replaced with ColXV-B were also designed to be used as controls. We next seeded isolated GFP^+^ motor neurons from 24 hpf *mnx1:gfp* embryos onto the micropatterns for 24 hrs to allow neurites to grow (Fig. 3C). The growth of GFP^+^-neurites was then quantified as the total number of pixels in ColXV-B and TnC areas (Fig. 3D). The results show that neurites are significantly more developed in the ColXV-B central route than in the TnC borders, whereas no difference was observed when motor neurons were seeded onto ColXV-B control micropatterns (Fig. 3D). Our finding uniquely unveils the importance of ECM cues in the vicinity of growing axons to locally instruct axon growth.

### ColXV-B is a molecular organizer that confers specific architecture to TnC

ColXV-B was reported to act as an ECM organizer (Hurskainen et al, 2005; Rasi et al, 2010). We thus assumed that ColXV-B could act as a molecular organizer of the motor path specialized ECM. To support this assumption, we first examined in details the temporal expression pattern of *tnc*, *col18a1* and *col15a1b* in the less differentiated cells of cluster 11 identified based on *smyhc1* expression level (Fig. 4A). We identified *col15a1b* as the earliest gene expressed by differentiating SMPs within the cluster 11, followed by *col18a1* and then *tnc* (Fig. 4A, Supplementary Table 2) while *col19a1* starts to be expressed in more mature SMPs in cluster 5 (Supplementary Table 2). Only the expression of *col15a1b* and *tnc* persists in the most differentiated SMPs in cluster 1 (Supplementary Table 4).

**Figure 4.**
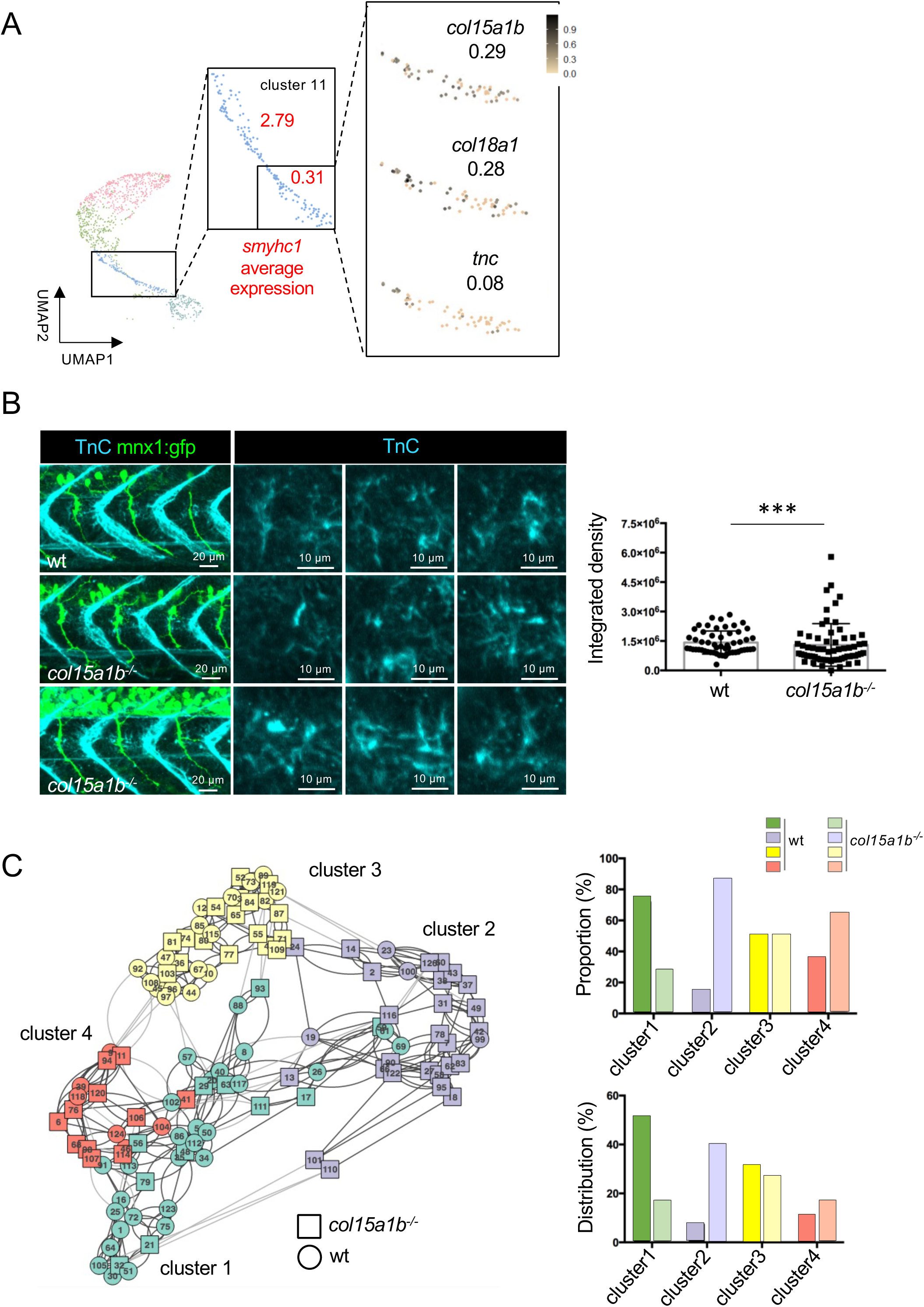
Lack of ColXV-B compromises TnC channel-like organization in the motor axon path. (**A**) Isolation of the cells with an average expression of *smyhc1* < 1 in the cluster 11 and UMAP feature plots of *col15a1b, col18a1 and tnc* expression patterns; values correspond to average expression. **(B)** Whole-mount immunostaining of 27 hpf *mnx1:gfp* and *col15a1b^-/-^;mnx1:gfp* embryos with anti-TnC (cyan) and anti-GFP (green, motor axons) antibodies. Zoomed image of TnC staining (left panel). Quantification (right panel). Statistical analysis was performed using a Kolmogorov-Smirnov test. ***p<0.0005. n^wt^ = 55, n*^col15a1b-/-^* = 65. Error bars are SEM. **(C)** Clustering analysis of 120 images of TnC staining along the motor path (left panel). One color represents one cluster. wt embryos (circle), *col15a1b^-/-^* embryos (square). Proportion and distribution of wt and *col15a1b^-/-^* mutant embryos in each cluster (right panel). For a same color, dark color represents wt embryos and light color represents *col15a1b^-/-^*embryos.

In order to determine whether ColXV-B acts as a molecular organizer of the ECM present along the motor path, the architecture of TnC deposits was analyzed in 27 hpf *col15a1b^-/-^* mutant embryos. Strikingly, lack of ColXV-B results in the apparent loss of the TnC characteristic organisation in the motor path along with a significant variability in TnC fluorescence intensity (Fig. 4B, and quantification). To verify if the channel-like organization of TnC in the motor path is lost in the absence of ColXV-B, an unsupervised clustering analysis was conducted on 120 images to group images with a similar distribution of TnC along the motor path (Fig. 4C, interactive visualization of the cluster partition at https://igfl.gitbiopages.ens-lyon.fr/ruggiero/zebrafishtnc/).

The distribution profile features a characteristic bimodal distribution shape when TnC exhibits a typical channel-like organization as in most images of wildtype compared to *col15a1b*^-/-^ embryos (Supplementary Fig. 4A). All analyses were blind to embryo types to avoid bias. Clustering analysis reveals four clusters when only two clusters are expected (Fig. 4C). Cluster 4 (red) comprises few images among them some display a curved channel shape. Cluster 3 (yellow) is equally composed of images of wt and mutant embryos for which TnC deposits display no specific distribution. This cluster mainly contains images that were a posteriori found to be highly pixelated (Supplementary Fig. 4B,C). However, cluster 2 (mauve) contains 88% images of *col15a1b^-/-^* mutant embryos that were shown a posteriori to exhibit no specific distribution whereas cluster 1 (green) consists in 73% images of wt embryos with a bimodal shape of TnC distribution.

Together with its earliest expression in differentiating SMPs (Fig. 4A), these results suggest that ColXV-B acts as a specific molecular organizer of TnC deposits in the motor path. Following this line of reasoning, we assumed that the absence of TnC expression does not affect ColXV-B organization in the motor path. We thus generated and characterized a *tnc* knock-out line using CrispR/Cas9 method to ascertain our assumption (Supplementary Fig. 5A-B). Interestingly, no difference in ColXV-B deposition was observed in the absence of TnC (Supplementary Fig. 5C). To conclude, SMPs leave behind ECM guidance cues that are essential for shaping the multiple ECM cues into a functional network that will locally guide motor axons as they exit from the spinal cord.

### ColXV-B influences motor axon navigation independently of TnC

As the lack of ColXV-B results in TnC desorganization in the motor path, we then asked whether the *col15a1b^-/-^* phenotype reported in (Guillon et al, 2016) is an indirect consequence of the loss of the characteristic TnC topology in the motor path. We reasoned that if it is the case, the motor axon phenotype of the *tnc*^-/-^ embryos should be identical to the *col15a1b^-/-^*phenotype. Knock-down of TnC was reported to result in excessive axon branching (Schweitzer et al, 2005). We thus analysed the motor axon phenotype of our newly generated *tnc^-/-^* line (no TnC, Supplementary Fig. 5B) and compared the potential defects in axon navigation to the ones observed in the *col15a1b^-/-^* mutants (loss of TnC architecture). We thus first quantified motor axon branching in wt, *tnc^-/-^* and *col15a1b^-/-^* embryos using “filament tracer” of Imaris software (Fig. 5A). Measurements of the total length of axon branches normalized to axon length signify that both *tnc^-/-^* and *col15a1b^-/-^* embryos abnormally develop excessive axon branches, even though the phenotype was significantly aggravated in *col15a1b^-/-^* embryos (Fig. 5B). Contrariwise, absence of TnC has no significant effect on motor axon length (4.8% increase compared to wildtype control, p-value >0.5) while *col15a1b^-/-^*embryos display a significant decrease in motor axon length (14.3% compared to wildtype control, p-value <0.0001), as we previously reported (Guillon et al, 2016). Together, these data strongly suggest that ColXV-B can influence axonal navigation through both TnC-dependent and TnC-independent mechanisms.

**Figure 5.**
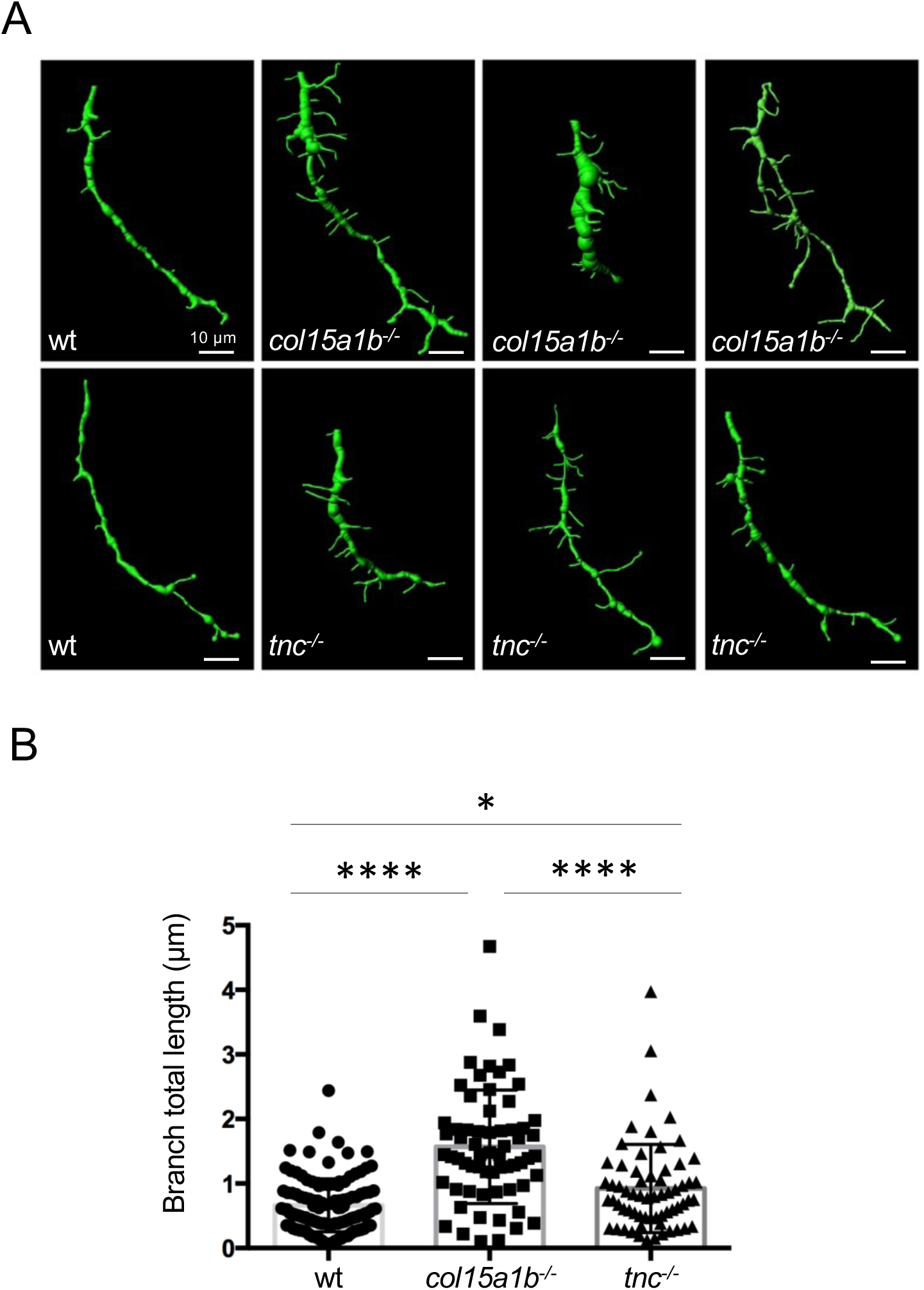
c*ol15a1b^-/-^*and *tnc^-/-^* embryos both display abnormal extra-branching phenotype. (**A**) 3D representation of motor axons using filament tracer (Imaris software) in wt, *tnc^-/-^* and *col15a1b^-/-^* mutant embryos. **(B)** Quantification of the cumulative length of all branches in wt, *col15a1b^-/-^* and *tnc^-/-^* mutant embryos. Statistical analysis was performed using a Kruskal-Wallis test. *p<0.05, ****p<0.0001. n^wt^ = 119, n*^col15a1b-/-^* = 64, n*^tnc-/-^*= 64. Error bars are SEM.

### ColXV-B deposits adjust motor axon microenvironment stiffness critical for outgrowth

ColXV-B is a hybrid collagen/proteoglycan molecule as it carries glycosaminoglycan (GAG) chains, that are predominantly chondroitin sulfate (CS) chains in mammals (reviewed in Bretaud et al, 2020) and in zebrafish as shown in Supplementary Fig. 6. The highly negatively charged GAG chains allow proteoglycans to sequester water that confers to tissues specific mechanical properties. There is growing evidence that axon elongation is influenced by tissue mechanics albeit differences in the response to the stiffness are observed depending on the type of neurons (reviewed in Athamneh et al, 2015). To investigate the impact of microenvironment stiffness in motor axon navigation, we prepared polyacrylamide hydrogels with different degrees of stiffness ranging from 5kPa to 40kPa. Hydrogels were coated with laminin, a well-known potent attractive ECM cue, to allow the motor neuron neurites to grow. Isolated 27 hpf *mnx1:gfp* motor neurons were cultivated for 24 hrs and neurite growth was imaged and measured as the total length of master segments per cell and the number of branches per cell using Fiji software (Fig. 6A, Supplementary Fig. 7). Quantification significantly shows that motor neurons develop more neurites when seeded onto soft than stiff hydrogels (5kPa vs 40kPa; Fig. 6B) indicative of a possible mechanism of action for ColXV-B by providing growing motor axons with a permissive soft microenvironment. To test this assumption, we next examined if *in vivo* ColXV-B deposits act on motor axon navigation by influencing the mechanical property in the vicinity of the motor axon path. We used atomic force microscopy (AFM) to measure the stiffness (elastic modulus) of the motor axon path on cross sections of 27 hpf *mnx1:gfp* and *col15a1^-/-^;mnx1:gfp* trunks. The results show that the motor axon path is stiffer in absence of ColXV-B (mean*^col15a1b-/-^* = 447.5 kPa) regarding to wt (mean^wt^ = 151.92 kPa) (Fig. 6C). Noticeably, injection of *shh* mRNA that increases deposition of ColXV-B (Guillon et al, 2016; Supplementary Fig. 3) leads to a significant decrease of the motor path stiffness (mean*^shh^*^-injected^ = 106.67 kPa) compared to wt (Fig. 6C). Our findings importantly extend the current belief that mechanosensing is critical for motor axon growth independently of chemical signals. In support to this, we have unravelled a mechanical role for ColXV-B in providing motor axons with a permissive soft ECM motor path.

**Figure 6.**
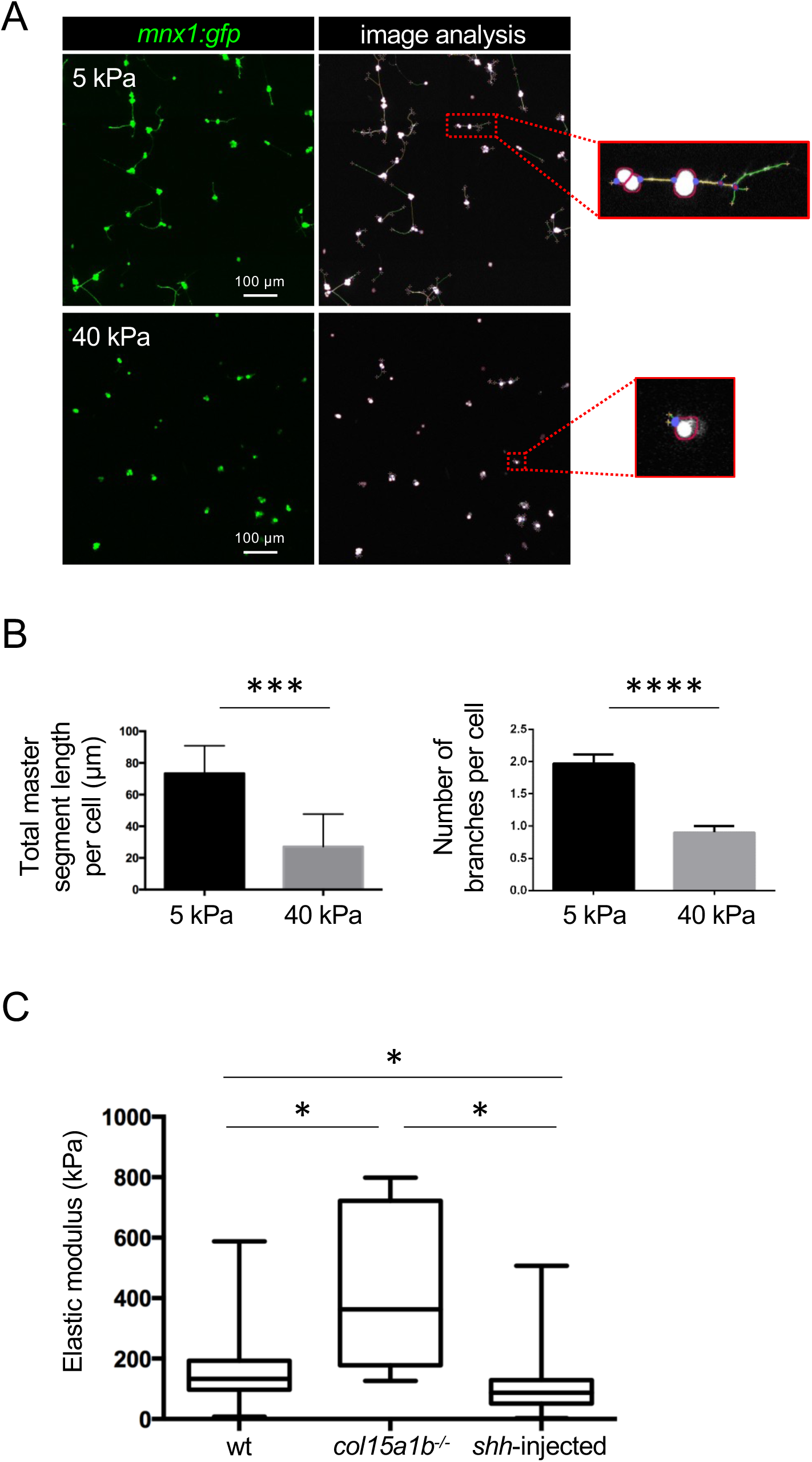
ColXV-B locally creates a soft microenvironment that elicits motor axon growth. (**A**) Confocal images of 27 hpf *mnx1:gfp* motor neurons cultivated on 5 or 40 kPa laminin-coated polyacrylamide hydrogels for 24 hours. Image analysis using ImageJ software shows motor neuron cell bodies (white), segments (yellow), and branches (green); zoomed images of the boxed areas on the right. **(B)** Quantification of the total master segment length and the number of branches per cell 24 hours after motor neurons seeding (n = 418 cells for 40 kPa gels; n = 803 cells for 5 kPa gels). Means of triplicates from 3 independent experiments are represented. Statistical analysis was performed using a Student’s t-test. Error bars are the SEM. ***p<0.0005; ****p<0.0001. **(C)** Elastic modulus in the motor axon path obtained from cryosections of 27 hpf wt, *col15a1b^-/-^*and *shh*-injected embryos in *mnx1:gfp* background. Statistical analysis was performed using Kruskal-Wallis and pairwise-Wilcoxon tests. *p<0.05. n^wt^ = 1342, n*^col15a1b-/-^* = 690, n*^shh^*^-injected^ = 400. Error bars are SEM.

## Discussion

ECM contact-mediated cues locally influence motor axon navigation and pathfinding. This is generally well-admitted (Letourneau, 2009; Stanic et al, 2016). However, the still open question is what are the ECM constituents of the motor path and how they assemble to provide an extrinsic axonal-promoting environment. We provide here compelling evidence that differentiating SMPs represent a major local source of ECM guidance cues, repellents and/or attractants whose dual function is reflected by a dual topology tailored to influence motor axon shape and growth during development.

Classic *in situ* hybridizations previously showed that developing SMPs (a.k.a adaxial cells) expressed ECM genes as *col15a1b* which expression was already detected in 13 hpf embryos (Bretaud et al, 2011) while *tnc* and *col19a1* transcripts were respectively detected in 16 hpf (Schweitzer et al, 2005) and 19hpf (Hilario et al, 2010) embryos. Using scRNAseq, we characterized the repertoire of ECM genes expressed by differentiating SMP and their temporal expression. A total of 36 core-matrisome genes are expressed by differentiating SMPs in both an overlapping and sequential pattern, providing ample evidence that SMPs are major local producers of the motor path ECM components. Importantly, our study assigns a specific gene signature to the principal morphological changes that SMPs undergo during their differentiation and hence sheds light on the understanding of the SMP lineage in zebrafish.

Our STRING analysis of SMP matrisomes reveal that most of the core-matrisome gene products interact to form a specialized BM containing both interlocked typical BM proteins and specific BM-associated proteins, namely TnC, ColXVIII, ColXIX and ColXV-B. While BM are built with major components found in all BM (a.k.a. the BM toolkit), they can display more complex composition and distinct physical properties that serve tissue-specific functions (reviewed in Sherwood, 2021). Interestingly, TnC, ColXVIII, ColXIX and ColXV-B have all been reported to influence motor axon navigation in zebrafish during axon development (Schweitzer et al, 2005; Schneider and Granato, 2006; Hilario et al 2010; Guillon et al, 2016), although only TnC (Schweitzer et al, 2005) and ColXV-B (Guillon et al, 2016) proteins were reported to be physically present in the axonal microenvironment. Using double immunostaining and loss of function studies, we marked an important step forward by showing that ColXV-B and TnC organize in a unique dual-topology that ideally serve axon navigation during development. These findings were further validated and corroborated by using motor path-like bioprinted micropatterns that demonstrate that the dual-topology of the ColXV-B and TnC immobilized cues is not only necessary but sufficient to control neurite navigation. Specialized BM have been described in several tissues including in different areas of the same cell, such as the myotendinous junction and the neuromuscular junction in myofibers (Sanes, 2003). Together, our data spot the motor path as a novel specialized BM whose composition and topology is tailored to locally guide motor axon towards their muscle target.

Tnc and ColXV-B deposition in the motor path by SMPs both depend on MusK signaling (Schweitzer et al, 2005; Guillon et al, 2016). However, our scRNA seq identifies *col15a1b* as the earliest BM-associated gene expressed in differentiated SMPs. Along this line, we showed that lack of ColXV-B leads to impaired TnC organization in the motor path. Conversely absence of TnC has no effect ColXV-B organization. This is in line with previous findings suggesting that human Collagen XV acts as an ECM organizer in cell assays and in tissue (Hurskaintern et al, 2010, Rasi et al, 2010). As such, the absence of ColXV-B indirectly results in a significant increase in axon branching, a phenotype described in TnC morphants (Schweitzer et al, 2005) and confirmed in our study using our *tnc* KO line. Likewise, old study showed that neurites exiting from chick spinal cord explants grow in straight paths with rare branch additions when plated onto TnC substrate (Wehrle and Chiquet, 1990). Together, these data suggest that, in addition to its role in axon guidance, ColXV-B may orchestrate the organization of SMP-derived ECM proteins to make the ECM motor path fully functional.

There is compelling evidence that growing neurons respond not only to chemical but also to mechanical signals. Axons can sense and respond to the mechanical property of their close environment (Koser, 2016; Martinez et al, 2021). The mechanisms underlying durotaxis are not fully elucidated. It is however well recognized that neurons are able to sense variation in their microenvironment stiffness through receptors localized at the growth cone (reviewed in Athamneh et al, 2015). Shortage of the mechanosensitive ion channel Piezo1 provoked axon pathfinding errors in *Xenopus* retinal ganglion cells (Koser et al, 2016). Interestingly, we showed that zebrafish motor axons are mechanosensitive and favourably extend neurites on soft substrate. We have shown that ColXV-B carries chondroitin sulfate chains, as its human and mouse counterpart (reviewed in Bretaud et al, 2020). Proteoglycans contribute to tissue viscoelasticity through their negatively charged sulfated glycosaminoglycans that are capable to bind and retain water in tissues (Murienne et al, 2015). *In vivo* application of chondroitin sulfate chains to xenopus brain decreased its stiffness and subsequently affected the trajectory of RGC axons (Koser et al, 2016). More specifically, lack of Collagen XV in mice resulted in a significant increase of the cardiac ECM stiffness (Rasi et al, 2010). Accordingly, our data strongly suggest that the presence of chondroitin sulfate chains endows ColXV-B with unique mechanical properties, locally perceived as an instructive signal for motor axon growth.

In conclusion, we demonstrate the central role of myotomal ColXV-B in the elaboration of a specialized BM with unique topology and mechanical properties that are locally critical for motor axon shape and navigation, corroborating the assumption that *COL15A1* should be viewed as a candidate gene for neuromuscular disorders and peripheral neuropathies with unresolved genetic cause. Importantly, our study not only provides fundamental insight into how ECM temporally and spatially orchestrates axonogenesis in vertebrates but will also help in the development of biomimetic ECM micropatterned scaffold designed to promote peripheral nerve regeneration.

## Materials and Methods

### Fish strain and maintenance

Zebrafish (AB/TU) maintenance and embryo collection were performed at the zebrafish PRECI facility (Plateau de Recherche Expérimentale de Criblage In vivo, UMS CNRS 3444 Lyon Biosciences, Gerland) in compliance with French government guidelines. Embryos obtained from natural spawning were raised following standard conditions. Developmental stages are given in hours post-fertilization (hpf) at 28.5°C according to morphological criteria (Kimmel et al, 1995). The *col15a1b* mutant strain carrying a nonsense mutation (*col15a1b*^sa12573^ allele) was obtained from the European Zebrafish Resource Center (Eggenstein-Leopoldshafen, Germany). Fish genotyping was performed as in Guillon et al, 2016. *tnc^2stopex3^* mutant zebrafish line was generated using CRISPR/Cas9 mediated genome editing according to Gagnon et al, 2014. A stop codon cassette was introduced leading to a premature termination codon in exon 3 (See Supplementary experimental procedure). Both mutant lines were outcrossed with Tg(*mnx1:gfp*) line to visualize motor neurons. All procedures have been approved by the local ethical committee and the ministry of higher education and research (APAFIS#27917-2020080513436447v7, APAFIS #37965-2022051900255627 v4)

### Recombinant proteins and antibodies against the NC10 domain of ColXV-B

Murine TnC, the zebrafish ColXV-B and ColXV-B NC1 domains (rNC10) were recombinantly produced in HEK-293 cells and purified as described in Supplementary Material and Methods. To generate guinea-pig antibodies to the zebrafish ColXV-B, strep tag was removed from the eluted proteins by thrombin (Sigma) digestion overnight at room temperature. ColXV-B NC1 domain was concentrated with spin-X concentrator (Corning) in PBS before its use. The purified domain was used to immunize guinea pigs and for affinity-chromatography purification of the immune sera (Covalab, Lyon, France). The generated anti-rNC10 antibodies were analysed for their specificity with whole-mount immunofluorescence staining of 27hpf embryos (Supplementary Fig. 2B).

### Chondroitin chains analysis

Digestion with Chondroitinase ABC of recombinant ColXV-B followed by Western blot analysis of the digested products and In Cell ELISA were used to analyse the presence of chondroitin sulfate on ColXV-B molecule as detailed in Supplementary Experimental Procedure.

### Fluorescence-activated cell sorting (FACS) of slow muscle progenitors

One cell stage AB/TU embryos were injected with 75 pg of s*myhc1:gfp* plasmid (gift from Stone Elworthy, University of Sheffield, Sheffield, United Kingdom) previously digested with meganuclease ISceI (Biolabs) in order to label slow muscle cells. GFP^+^ embryos were dissected at 18 hpf (n=200) by removing the head and the yolk sac under anaesthesia (tricaine 0.016%). Trunks were placed in 1 ml 0.1X MMR solution (Marc’s modified Ringers, 100 mM NaCl, 2 mM KCl, 2 mM CaCl2, 1mM MgSO4, 5 mM hepes, 0.1 mM EDTA) at 4°C until the end of the dissection to preserve the tissue. Trunks were then dissociated in trypsin dissociation buffer (115 mM NaCl, 2.5 mM KCl, 8 mM hepes, 0.4 mM EDTA, 0.05% trypsin, 0.0165µg/µl DNase) at 37°C (350 rpm, 1 hour) in a thermomixer. One mL medium (80% Leibovitz medium, Life Technologies, 10% 0.1X MMR, 10% fetal bovine serum, FBS) was then added to the cell suspension to inhibit the trypsin. Cells were filtered using a 70 µm nylon strainer (BD Falcon) and centrifugated (1200 rpm, 10 min, 4°C). The cell pellet was resuspended in 400 µl PBS 1% FBS, 1 mM EDTA. GFP^+^ cells were selected by FACS (FACSCANTO, ANIRA-cytometry, UMS 3444, SFR Biosciences Gerland). 20000 cells were collected in 500 µl PBS 10% FBS and immediately used for scRNAseq.

### Single-cell RNA sequencing analysis and bioinfomatics

ScRNAseq was performed using 10X Genomics technology and Chromium Next GEM Single Cell 3’ Reagent Kits v3.1 kit. GFP^+^ FACS-sorted cells were centrifugated (1200 rpm, 10 min, 4°C) to concentrate at 900 cells/µl (cell concentration estimated using a Malassez cell, C-Chip, NanoEnTek). 5000 cells were loaded into the Chromium Controller and the library constructed following manufacturer’s instructions. The concentration of the library was assessed by Qubit HS DNA quantification and its quality evaluated by Tapestation 4150 analysis with HS D5000 screentape. The single-indexed (SI-GA-D2) library was sequenced on a High Output run on an Illumina NextSeq500 sequencer with 28 cycles for R1 and 132 cycles for R2. The run generated more than 600M PF reads. ScRNAseq data was processed into transcript count tables using Cell Ranger software v6.0.1 developed by 10X Genomics. Raw base call (bcl) files generated by Illumina sequencer were converted into FASTQ files using the *cellranger mkfastq* pipeline and default parameters generating 608,065,849 reads. FASTQ files were then processed using the *cellranger count* pipeline to perform a firstalignment against the zebrafish genome (GRCz11, Ensembl GCA_000002035.4), filtering, barcode counting and UMI counting. The gfp gene was added to this genome to perform a second mapping. For the first mapping,we finally recovered 4832 cells with a median UMI counts per cell of 30047 and a median genes per cell of 3700. For the second mapping, we recovered 4998 cells with a median UMI counts per cell of 30808 and a median genes per cell of 3737. Barcodes.tsv.gz, features.tsv.gz and matrix.tsv.gz files obtained after both mappings were then imported to R software to be processed. Seurat package v4.3.0 was used for quality control following criteria as below: nFeature > 800 and nFeature < 6500; nCount > 5000 and nCount < 100000; percent.mt < 3, data filtration, dimensional reduction, graph-based clustering, and identification of signature genes. The gene expression threshold was set to 0.25. Raw reads for this project were deposited to SRA database under BioProject accession N°xxxxx (pending, submission number, SUB13190716).

### Immunostaining

Whole-mount immunostainings of 27 hpf embryos were performed as previously described (Guillon et al, 2016). For Sonic Hedgehog (Shh) signaling-forced activation, one-cell stage wildtype embryos were injected with 75 pg of *shh* mRNA and immunostaining was then performed on whole-mount or cryosections of 27 hpf *shh*-injected embryos as previously described (Guillon et al, 2016). Primary and secondary antibodies were used at indicated dilutions: guinea pig polyclonal anti-ColXV-B (see Supplementary Experimental Procedures) (1:1500); rabbit anti-Tenascin-C (1:500, USBiological); mouse monoclonal F59 antibody (1:10, Developmental Studies Hybridoma Bank [DSHB]); chicken anti-GFP (1:1000, Abcam); goat anti-mouse or anti-rabbit or anti-guinea pig IgG coupled to AlexaFluor-488 or AlexaFluor-546 or AlexaFluor-647 (1:500, Invitrogen), anti-chicken IgG coupled to AlexaFluor-488 (1:1500, Invitrogen). Embryos were observed and imaged at the level of the yolk sac extension using an inverted confocal microscope (Zeiss LSM 780). Images are 3D projections using Imaris software or Image J *Z* projections of confocal *Z* stacks as mentioned in the figure legends. Images were acquired with identical confocal settings and were equally post-treated with ImageJ software to value differences in signal intensity.

### Image analysis of Tenascin-C deposits and quantification

For each embryo (n = 11 for wt and n = 13 for *col15a1b^-/-^*), regions of interest covering one somite were individually defined using post processing image create subset of Zen software (Zeiss) from 5 adjacent somites. Individual images were randomly renamed to work as blind. Background for TnC staining was removed on Imaris software using clipping plane to keep only TnC staining in the motor path and images of TnC staining were created using snapshot with or without motor neuron staining. One image saturated for TnC staining was also created to better select on Image J the area of the presence of TnC. TnC staining images were treated in Image J using a macro. Saturated images were transformed into 8-bit images and phansalkar auto local threshold was applied to create a binary image. After TnC staining manual removing in the vertical myosepta, TnC staining in the motor path was selected and used as a mask to apply on the original (non-saturated) 8-bit image to remove the outside motor path staining. Finally, integrated density of TnC staining in the motor path was measured.

### TnC channel organization analysis

An unsupervised clustering analysis was conducted to discriminate the TnC channel organization between wt and *col15a1b^-/-^* mutant embryos. First, each 8-bit image was truncated horizontally (x-axis) around the channel center to focus on a common area of interest: the motor path. Following the truncation, the TnC distribution along the motor path of each image was estimated by vertically (along the y-axis) summing non-zero pixels. The resulting distribution curves were smoothed by a Gaussian filter to remove spurious noise and normalized. A dissimilarity matrix was then built by evaluating the Hellinger distance, between each pair of curves. Finally, a graph partition was constructed from the dissimilarity matrix with the Leiden algorithm to cluster together similar 8-bit images. Mathematical background of each step is detailed in the Supplementary Material.

### Motor axon analysis

In all embryos acquired, regions of interest containing the motor axon were individually defined using post processing image create subset of Zen software (Zeiss) from 5 adjacent somites. Individual images were randomly renamed to work as blind. Filament tracer from Imaris software (Bitplane) was used to measure axon branching (collaterals and terminal arborization) in individual somite. Automatic detection was used to track the axon and the seed points threshold was adjusted to detect either the axon or the axon with the branches. The total length of the axon was subtracted from the total length of axon with branches to determine the total length of only branches. The data were then normalized with the length of the axon measured on Fiji.

### Double micropatterning

Round coverslips of 32 mm diameter were cleaned with isopropanol and activated with oxygen plasma at 0.4 mbar with a power of 100% for 2 minutes. The treated surface was coated with poly-D-lysine solution (PLL, Thermofisher Scientific, 100 µg/ml) for 30 minutes and then rinse with deionized water and Hepes buffer (0.1M pH 8.3). PLL-treated coverslips were then passivated with mPEG-SVA solution (methoxypolyethylene glycol-succinimidyl valerate; Laysan Bio; 100 mg/ml in 0.1M Hepes buffer pH 8.3) for 1 hour, rinsed with deionized water and air dried to be stored at 4°C in the dark for few days.

For patterning, a PRIMO system (Alveole Lab) mounted on an epifluorescence microscope (Nikon Eclipse Ti2) was used. Patterns consisting of outside lines of 8 µm thickness and inner lines of 15 to 25 µm thickness on 70 µm of length (total of repeated 4000) were drawn using Inkscape software. A passivated coverslip was placed into a sample holder, covered with photoactivator mix (50 µl 70% ethanol, 1 µl surfactant, 5 µl PLPP gel, Alveole Lab) and mounted on the microscope (20X magnification) to let it dry. Using Leonardo software (Alveole Lab), a circular ROI (32 mm diameter) was aligned with the coverslip. The focus was adjusted with PFS and the first pattern was illuminated at 30 mJ/mm^2^ with a laser power of 100%. The coverslip was rinsed four times with deionized water. 1 mL of PLL (100 µg/ml) was first incubated for 20 min to improve the TnC coating. 400 µl of solution of recombinant TnC (Supplementary Materials and Methods) or purified human TnC (Merck Millipore) (15 µg/ml in 0.01% PBS-Tween) were then incubated for 30 min. The coverslip was then blocked with 1% bovine serum albumin (BSA) in deionized water for 30 minutes. A photoactivator mix was prepared as previously with 50% ethanol and spread on the coverslip. After complete drying, the second pattern was aligned on the first one and illuminated with the same parameters. The holder was removed from the microscope and the coverslip was incubated with 500 µl of zebrafish ColXV-B enriched medium from HEK-293 cells for 20 min. Coverslips can be stored at 4°C in deionized water for few days. After seeding FACS-isolated motor neuron on micropatterns and cultivating them for 24 hrs at 28°C, GFP fluorescence was acquired on inverted Zeiss LSM850 confocal. The signal was also acquired with the 561nm laser and the display was strongly increased for the analysis to localize micropatterns. Fiji macro was developed to detect the presence of the neurite in the 3 areas covered with TnC and ColXV-B based on the number of pixel/µm^2^. The data were then expressed in % of the total pattern area.

### FACS isolation of motor neurons

24 hpf *mnx1:gfp* zebrafish embryos were used to isolate motor neurons following the same protocol as for slow muscle cells. GFP^+^ cells were selected by FACS (FACSCANTO, ANIRA-cytometry, UMS 3444, SFR Biosciences Gerland) and directly collected in 100 µl neurotrophic factor enriched medium (50% Leibovitz medium, 49% 0.1X MMR, 1% FBS, BDNF 1:10000, GDNF 1:10000, CNTF 1:1000, Zellshield 1:200). The cell concentration was estimated using a Malassez cell (C-Chip, NanoEnTek) and 9000 cells were seeded on coverslips covered with PAA hydrogels of different stiffness in 24 well plates.

### Polyacrylamide hydrogel substrates

12 mm round silanized coverslips (treatment with 0.1% acetic acid, 0.1% bind silane in methanol) were used as a final support for the polyacrylamide (PAA) hydrogels. 20×20 mm square coverslips were coated with laminin (Sigma, 20 µg/ml in 100 mM sodium bicarbonate) on parafilm for 30 minutes. PAA hydrogels of 5 and 40 kPa were prepared as previously described (Tse and Engler, 2010) and a droplet of 5 µl was applied on the laminin-coated coverslip to obtain a hydrogel of 30 µm thickness. The silanized coverslip was then deposited on the PAA droplet. After polymerisation, PAA hydrogels were rehydrated with water and detached from square coverslip using a scalpel. PAA hydrogels were stored in water at 4°C in 24 well plates and placed at 28°C two hours before cell seeding.

### Neurite tracing and image analysis

FACS-sorted motor neurons were seeded on PAA gels for 24 hrs at 28°C and live-imaged on Zeiss confocal microscope (LSM780). One field from 3 PAA gels for each stiffness condition was acquired. Quantification of cell bodies, segments and branches on 1.1 x 1.2 mm confocal images of motor neurons was performed using Fiji macro (n = 3 experiments with triplicates).

### Atomic force microscopy analysis

After fixation of 27 hpf embryos in 4% paraformaldehyde, 4°C overnight, transverse cryosections were performed as previously (Guillon et al, 2016) on Tg(*mnx1:gfp*) injected or not with *shh* mRNA (100 pg) at 1 cell-stage and on homozygous *col15a1b^sa12573^;mnx1:gfp*. Elastic modulus of the environment of the motor axon was performed on selected sections where the axon was visible. Experiment was carried out by Biomeca (Lyon, France) using an atomic force microscope (Bioscope Catalyst-Bruker®) mounted on a fluorescent macroscope (MacroFluo, Leica). 300 measurements were performed on a ROI of 15µm^2^.

### Statistical analysis

Statistical analyses were carried out using GraphPad Prism. Depending on the experiments, the statistical methods to compare datasets were different and include Shapiro-Wilk test, Student’s t-test, Kruskal-Wallis test, pairwise-Wilcoxon test, Kolmogorov-Smirnov test and one way ANOVA as indicated in the legend of the figures.

## Acknowledgments

We are grateful to Dr Gertraud Orend (Univ Strasbourg, France) for her helpful advice in handling Tenascin C. We thank Chloé Exbrayat-Héritier for her technical assistance in embryo dissection, Guillaume Marcy (University of Lyon), Nathan Gil and Camille Guillermin (IGFL, ENS de Lyon) for their help with single cell analysis and Vladimir Misiak (LIPhy, Univ Grenoble) for his assistance in the fabrication of the micropatterns. We acknowledge the contribution of the Spatial-Cell-ID EquipEx+ facility for single cell analysis and of the SFR Biosciences (UAR3444/CNRS, US8/Inserm, ENS de Lyon, Université Claude Bernard Lyon 1, Lyon, France) facilities, notably the zebrafish facility (PRECI, Laure Bernard and Robert Renard) and the flow-cytometry facility (Sébastien Dussurgey) and Denis Ressnikoff at the CIQLE facility (University Lyon 1) for his help in image analysis (Imaris).

## Author contributions

Conceptualization: LNB, SB, FR; Methodology: LNB, MB, LG, BG, IS, GC, MJGM, EG, EV, YGH, SB; Resources: MK, YGH; Investigation: LNB, MB, LG, SB; Visualization: LNB, MB, EG, GC, SB; Supervision: FR, SB; Funding acquisition: FR; Writing—original draft: LNB; Writing— review & editing: LNB, SB, FR

## References

1. B. D. Ackley, J. R. Crew, H. Elamaa, T. Pihlajaniemi, C. J. Kuo, J. M. Kramer, The Nc1/Endostatin Domain of Caenorhabditis elegans Type Xviii Collagen Affects Cell Migration and Axon Guidance. Journal of Cell Biology. 152, 1219–1232 (2001).

2. A. I. M. Athamneh, D. M. Suter, Quantifying mechanical force in axonal growth and guidance. Front. Cell. Neurosci. 9 (2015).

3. S. Baxendale, C. Davison, C. Muxworthy, C. Wolff, P. W. Ingham, S. Roy, The B-cell maturation factor Blimp-1 specifies vertebrate slow-twitch muscle fiber identity in response to Hedgehog signaling. Nat Genet. 36, 88–93 (2004).

4. C. E. Beattie, Control of motor axon guidance in the zebrafish embryo. Brain Research Bulletin. 53, 489–500 (2000).

5. R. R. Bernhardt, M. Schachner, Chondroitin Sulfates Affect the Formation of the Segmental Motor Nerves in Zebrafish Embryos. Developmental Biology. 221, 206–219 (2000).

6. S. Bretaud, A. Pagnon-Minot, E. Guillon, F. Ruggiero, D. Le Guellec, Characterization of spatial and temporal expression pattern of *col15a1b* during zebrafish development. Gene Expression Patterns. 11, 129–134 (2011).

7. S. Bretaud, E. Guillon, S.-M. Karppinen, T. Pihlajaniemi, F. Ruggiero, Collagen XV, a multifaceted multiplexin present across tissues and species. Matrix Biology Plus. 6–7, 100023 (2020).

8. S. Bretaud, P. Nauroy, M. Malbouyres, F. Ruggiero, Fishing for collagen function: About development, regeneration and disease. Seminars in Cell & Developmental Biology. 89, 100– 108 (2019).

9. S. Carbonetto, M. Lindenbaum, The basement membrane at the neuromuscular junction: a synaptic mediatrix. Current Opinion in Neurobiology. 5, 596–605 (1995).

10. A. G. Clementz, M. J. Mutolo, S.-H. Leir, K. J. Morris, K. Kucybala, H. Harris, A. Harris, Collagen XV Inhibits Epithelial to Mesenchymal Transition in Pancreatic Adenocarcinoma Cells. PLoS ONE. 8, e72250 (2013).

11. D. F. Daggett, C. R. Domingo, P. D. Currie, S. L. Amacher, Control of morphogenetic cell movements in the early zebrafish myotome. Developmental Biology. 309, 169–179 (2007).

12. B. J. Dickson, Molecular Mechanisms of Axon Guidance. 298 (2002).

13. J. A. Gagnon, E. Valen, S. B. Thyme, P. Huang, L. Ahkmetova, A. Pauli, T. G. Montague, S. Zimmerman, C. Richter, A. F. Schier, Efficient Mutagenesis by Cas9 Protein-Mediated Oligonucleotide Insertion and Large-Scale Assessment of Single-Guide RNAs. PLoS ONE. 9, e98186 (2014).

14. E. Guillon, S. Bretaud, F. Ruggiero, Slow Muscle Precursors Lay Down a Collagen XV Matrix Fingerprint to Guide Motor Axon Navigation. J. Neurosci. 36, 2663–2676 (2016).

15. J. D. Hilario, C. Wang, C. E. Beattie, Collagen XIXa1 is crucial for motor axon navigation at intermediate targets. Development. 137, 4261–4269 (2010).

16. M. Hurskainen, L. Eklund, P. O. Hägg, M. Fruttiger, R. Sormunen, M. Ilves, T. Pihlajaniemi, Abnormal maturation of the retinal vasculature in type XVIII collagen/endostatin deficient mice and changes in retinal glial cells due to lack of collagen types XV and XVIII. FASEB j. 19, 1564–1566 (2005).

17. M. Hurskainen, F. Ruggiero, P. Hägg, T. Pihlajaniemi, P. Huhtala, Recombinant Human Collagen XV Regulates Cell Adhesion and Migration. Journal of Biological Chemistry. 285, 5258–5265 (2010).

18. C. B. Kimmel, W. W. Ballard, S. R. Kimmel, B. Ullmann, T. F. Schilling, Stages of embryonic development of the zebrafish. Dev. Dyn. 203, 253–310 (1995).

19. D. E. Koser, A. J. Thompson, S. K. Foster, A. Dwivedy, E. K. Pillai, G. K. Sheridan, H. Svoboda,

20. M. Viana, L. D. F. Costa, J. Guck, C. E. Holt, K. Franze, Mechanosensing is critical for axon growth in the developing brain. Nat Neurosci. 19, 1592–1598 (2016).

21. P. C. Letourneau, Axonal Pathfinding: Extracellular Matrix Role. (2009).

22. D. Li, C. C. Clark, J. C. Myers, Basement Membrane Zone Type XV Collagen Is a Disulfide-bonded Chondroitin Sulfate Proteoglycan in Human Tissues and Cultured Cells. Journal of Biological Chemistry. 275, 22339–22347 (2000).

23. G. F. Martínez, J. Fagetti, G. Vierci, M. M. Brauer, N. Unsain, A. Richeri, Extracellular matrix stiffness negatively affects axon elongation, growth cone area and F-actin levels in a collagen type I 3D culture. J Tissue Eng Regen Med. 16, 151–162 (2022).

24. E. Melançon, D. W. C. Liu, M. Westerfield, J. S. Eisen, Pathfinding by Identified Zebrafish Motoneurons in the Absence of Muscle Pioneers. J. Neurosci. 17, 7796–7804 (1997).

25. F. Meyer, B. Moussian, Drosophila multiplexin (Dmp) modulates motor axon pathfinding accuracy: Dmp modulates motor axon pathfinding. Development, Growth & Differentiation. 51, 483–498 (2009).

26. B. J. Murienne, J. L. Jefferys, H. A. Quigley, T. D. Nguyen, The effects of glycosaminoglycan degradation on the mechanical behavior of the posterior porcine sclera. Acta Biomaterialia. 12, 195–206 (2015).

27. P. Nauroy, S. Hughes, A. Naba, F. Ruggiero, The in-silico zebrafish matrisome: A new tool to study extracellular matrix gene and protein functions. Matrix Biology. 65, 5–13 (2018).

28. K. Rasi, J. Piuhola, M. Czabanka, R. Sormunen, M. Ilves, H. Leskinen, J. Rysä, R. Kerkelä, P. Janmey, R. Heljasvaara, K. Peuhkurinen, O. Vuolteenaho, H. Ruskoaho, P. Vajkoczy, T. Pihlajaniemi, L. Eklund, Collagen XV Is Necessary for Modeling of the Extracellular Matrix and Its Deficiency Predisposes to Cardiomyopathy. Circ Res. 107, 1241–1252 (2010).

29. S. Ricard-Blum, F. Ruggiero, The collagen superfamily: from the extracellular matrix to the cell membrane. Pathologie Biologie. 53, 430–442 (2005).

30. J. R. Sanes, The Basement Membrane/Basal Lamina of Skeletal Muscle. Journal of Biological Chemistry. 278, 12601–12604 (2003).

31. V. A. Schneider, M. Granato, The Myotomal diwanka (lh3) Glycosyltransferase and Type XVIII Collagen Are Critical for Motor Growth Cone Migration. Neuron. 50, 683–695 (2006).

32. J. Schweitzer, T. Becker, J. Lefebvre, M. Granato, M. Schachner, C. G. Becker, Tenascin-C is involved in motor axon outgrowth in the trunk of developing zebrafish. Dev. Dyn. 234, 550– 566 (2005).

33. D. R. Sherwood, Basement membrane remodeling guides cell migration and cell morphogenesis during development. Current Opinion in Cell Biology. 72, 19–27 (2021).

34. K. Stanic, N. Saldivia, B. Förstera, M. Torrejón, H. Montecinos, T. Caprile, Expression Patterns of Extracellular Matrix Proteins during Posterior Commissure Development. Front. Neuroanat. 10 (2016).

35. H. L. Stickney, M. J. F. Barresi, S. H. Devoto, Somite development in zebrafish. Dev. Dyn. 219, 287–303 (2000).

36. J. R. Tse, A. J. Engler, Preparation of Hydrogel Substrates with Tunable Mechanical Properties. Current Protocols in Cell Biology. 47 (2010).

37. T. Unsoeld, J.-O. Park, H. Hutter, Discoidin domain receptors guide axons along longitudinal tracts in C. elegans. Developmental Biology. 374, 142–152 (2013).

38. B. Wehrle, M. Chiquet, Tenascin is accumulated along developing peripheral nerves and allows neurite outgrowth in vitro.

39. J. Zhang, J. L. Lefebvre, S. Zhao, M. Granato, Zebrafish unplugged reveals a role for muscle-specific kinase homologs in axonal pathway choice. Nat Neurosci. 7, 1303–1309 (2004).

40. J. Zhang, M. Granato, The zebrafish *unplugged* gene controls motor axon pathway selection. Development. 127, 2099–2111 (2000).

