## Supplemental Material and Method – Fig S1-7 – Table 1-5 for "Dual-topology of collagen XV and tenascin C acts in concert to guide and shape developing motor axons"

### Supplementary Material and Methods

Recombinant proteins and antibodies against the NC1 domain of ColXV-B

The zebrafish ColXV-B NC1 domain (LK391962, aa: 969-1245) and the murine TNC (NP_035737.2, aa: 174-2019; cDNA gift by R. Chiquet-Ehrismann) were cloned into the PCEP4 expression vector (Invitrogen). A BM40 signal peptide and a N-terminal Twin tag was added to each cDNA and the final plasmids were verified by sequencing. To generate stable cell line, HEK293 EBNA cells were transfected with the expression vector using Fugene HD (Promega). 48 hrs after transfection, the medium was replaced with DMEM/F12 medium with 10% FCS containing 0.5 µg/ml puromycin (Sigma) for selection. After one week of selection, the cells were transferred to triple flasks (Thermo Fisher Scientific) and the conditioned media harvested every 2-3 days. The recombinant proteins were then purified from supernatants using the Streptactin matrix (IBA, Lifesciences) following the manufacturer's guidelines. The eluted protein was then dialyzed against PBS and stored at -70°C until use.

The ColXV-B was produced in HEK_293 cells as previously described (Guillon et al, 2016). A clonal selection was performed in DMEM/F12 medium with 10% FCS containing 250 µg/ml of hygromycine (Sigma) and the ColXV-B production yield of the different clones was tested with immunofluorescence staining and SDS-PAGE analysis. Clone 14 was selected and ColXV-B-enriched medium from clone 14 cells grown in 50 µg/ml of vitamin C and absence of FCS was collected after 24 hrs and stored at -70°C until use.

Mathematical analysis for the TnC distribution analysis

TnC staining values of an 8-bit image obtained from the Tn-C deposition assessment can be represented as a matrix $\mathrm{X}$ of shape $(1294,970)$ and we will denote by, $\mathrm{X(i,j)}, \mathrm{i} \in \{1,\dots,1294\}, \mathrm{j} \in \{1,\dots,970\}$, TnC staining value at the pixel with coordinates $(\mathrm{i},\mathrm{j})$.The TnC channel organization analysis was comprised of four steps: image truncation, TnC distribution estimation, construction of the dissimilarity matrix, construction of the clustering graph. Mathematical analysis of each step is detailed hereafter.

For image truncation, each 8-bit image was first truncated horizontally around the channel center. The $\mathrm{i}$-coordinate of the channel center (red lines in Fig. S3), denoted by $\mathrm{c_i}$, was selected manually for each image. The truncation radius was set equal to 75 pixels. Then, the matrix associated with the truncated image was defined as,

$X_T(i,j) = X(c_i - 75 + i, j),\ i \in \{1,\ldots,1294\}, j \in \{1,\ldots,970\}$. Hence, the truncated 8-bit images are 150 pixels width large.

For TnC distribution measurement, TnC distribution along the motor path was estimated for each truncated image by vertically summing (along the j-coordinate) non-zero pixels. The TnC distribution can be represented as an array $z = (z_1, z_2, \ldots)$ of length 150 such that,

$z_i = \sum_{j=1}^{970} H\left(X_T(i,j)\right), i \in \{1,\ldots,150\}$ ,

where $H$ is the Heaviside step function defined by:

$H(x) = 1_{x>0} = \begin{cases} 1 & \text{if } x > 0 \\ 0 & \text{if } x \leq 0 \end{cases}$ .

Each array $z$ can be plotted as a curve. Most of the obtained curves were noisy (presence of local spikes) due to sharp local variations in non-zero pixel counts. To remove this noise, each curve was smoothed by applying a 1D convolution filter with a Gaussian kernel. The idea behind this filtering step is to correct each value $z_i$ by a weighted sum of its neighboring values, where the weights are obtained from a Gaussian kernel. The filtered curve is defined by the array $\tilde{z}$:

$$\tilde{z}_i = \sum_{k=-L}^{L} z_{i-k} w_k \, , w_k = \frac{1}{\sqrt{2\pi}\sigma} \exp\left(\frac{-k^2}{2\sigma^2}\right),$$

where $w_k$ are the Gaussian weights, $2L$ the number of neighbouring values and $\sigma$ the standard deviation. The filtering step was performed with the gaussian_filter1d function from the Python package Scipy and with parameters $\sigma = 5$ and $L = 20$. Two examples of filtered curves are shown in Fig. S3 (blue curves).

For the construction of the dissimilarity matrix, a statistical distance, called Hellinger distance, was used to measure the dissimilarity between the TnC distributions of two images. The closer the distance to zero, the more similar the two distributions. Since the Hellinger distance requires probability distributions as inputs, the filtered arrays were first normalized,

$\tilde{z}_i = \frac{\tilde{z}_i}{\sum_i \tilde{z}_i}$,

then the discrete Hellinger distance was computed for each pair of filtered arrays,

$$d_H(\tilde{z}, \tilde{z}') = \frac{1}{\sqrt{2}} \sqrt{\sum_i \left(\sqrt{z_i} - \sqrt{z'_i}\right)^2} ,$$

and stored in the dissimilarity matrix $D$ of shape $(120{,}120)$.

For the construction of the clustering graph, a k-nearest neighbour graph was constructed from the dissimilarity matrix and infers the k-closest neighbours of each filtered array. The function

NearestNeighbors from the Pyton package scikit-learn was used to construct the k-neighbours graph with parameter $\mathrm{k}=5$.
An adjacency matrix was then computed from the k-neighbours graph and fed as input to the Leiden algorithm to detect a partition of clusters. The Leiden algorithm uses a parameter $\gamma \in [0,1]$, called resolution, which drives the number of clusters found. Higher resolutions lead to more clusters. The Python package leidenalg was used to apply the Leiden algorithm with a small resolution, $\gamma = 0.1$, as we knew we were looking for a small number of clusters.

Generation of *tnc* zebrafish mutant line
*Tnc* mutant zebrafish line was generated using CRISPR/Cas9 mediated genome editing. Gene specific guide RNA (sgRNA) (CCAGGATCAAGGGCGTTGTGAGG) was determined using CRISPOR (crispor.tefor.net) to target exon 3 and synthesized using EnGen sgRNA synthesis kit from New England Biolabs. A stop codon cassette was used as a matrix to create a premature termination codon as described in Gagnon et al, 2014. One cell-embryos of AB/TU background were injected with 500pg of sgRNA, 600 pg of Cas9 protein (NEB) and 3 µl of stop codon cassette. 2 F0 founders were identified by PCR using a primer in the stop codon cassette (5'-GGCGTTTAAACCTTAATTAAGC-3') and a primer in *tnc* exon 3 (5'-CGCAGTCTTCCCCAGTAAAG-3'). The fish were used to raise F1 fish by outcrossing with Tg(*mnx1 :gfp*)$^{ml2}$. After sequencing the F1 offspring, 5 fish carrying the same genotype corresponding to an insertion of 127 nucleotides (2 stop codon cassettes with the homology arms) were selected. F2 embryos from the incrossed heterozygote F1 fish were genotyped after immunofluorescence using 2 primers closed to the target sequence allowing to distinguish directly between heterozygote and homozygote (5'-GAGCCAGAATGCCCAAACTA -3' forward, 5'-CCCGTCGATACACTCTCCAT-3' reverse).

Chondroitin chains analysis
500 µl of ColXV-B enriched medium from HEK293 cells were digested with chondroitinase ABC (50 units, *proteus vulgaris,* Sigma) for 3 hrs at 37 °C in 50 mM Tris-HCl pH 8, 50 mM sodium acetate, 100 µg/ml BSA, 1 mM PMSF. Western blot analysis of the digested or non-digested media with NC10 antibodies was performed as previously (Guillon et al, 2016). HEK293 cells expressing ColXV-B and untransfected cells as control were used to perform In cell ELISA. Cells were seeded at 400,000 cells per well in 96 well plate pre-coated with poly-D-lysine (100µg/ml, Sigma). The following day, medium was replaced with fresh medium supplemented with vitamin C (50µg/ml, Sigma). After 24 hrs, cells were fixed with 4% PFA, permeabilized with

0.1% triton X-100 and saturated with 1% BSA. Then, cells were incubated overnight at 4°C with or without anti-C6S antibody (m473 HD, gift by Adreas Faissner, Germany) followed by incubation with anti-rat IgG-HRP conjugate and TMB substrate (BD Biosciences). After addition of 2M H2SO4, absorbance was measured at 595 nm with a microplate reader (Multiskan FC, Thermoscientific). Levels of chondroitin sulfate were measured to normalize to relative cell seeding density values obtained with Janus Green whole-cell stain (Abcam).

### Supplementary Figures

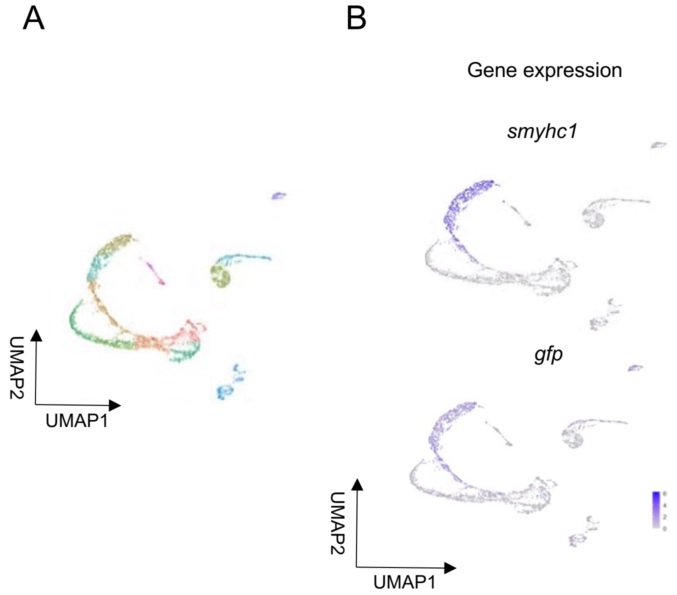


**Figure S1.** *smyhc1* positive cells express the *gfp*. (A) UMAP plot obtained after mapping on the zebrafish genome containing the *gfp* gene. (B) UMAP feature plots showing the expression patterns of *smyhc1* and *gfp*.

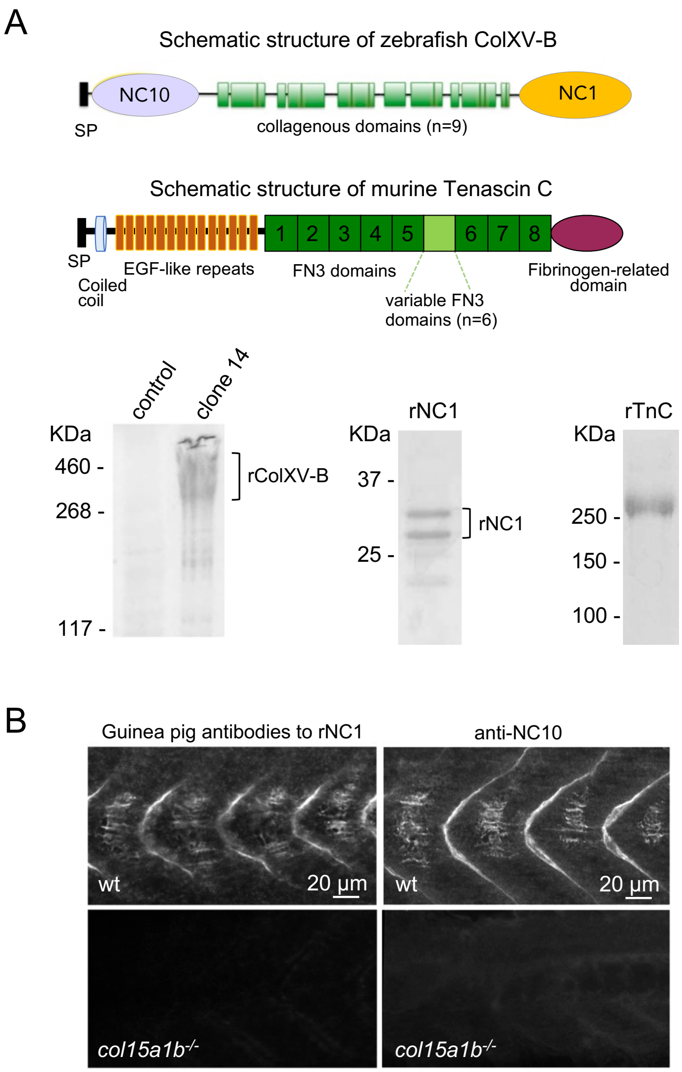


**Figure S2.** Recombinant ECM protein production and characterization of the guinea pig antibodies against zebrafish ColXV-B NC1 domain. (A) Schematic structure of ColXV-B and TnC (upper panel). NC, non-collagenous domains, numbered from the C-terminus. SP, signal peptide; proteins are not drawn to scale. Full-length ColXV-B and ColXV-B NC1 domain and TnC were recombinantly produced in HEK293 cells. Bottom panel, from left to right: SDS-PAGE analysis of 1 ml of clone 14 (rColXV-B) conditioned medium and untransfected HEK293 medium (control) (6% acrylamide); of 2µg recombinant ColXV-B NC1 domain (rNC1) (10% acrylamide) and 2µg recombinant Tenascin C (rTnC) (6% acrylamide). Molecular weight markers on the left side. (B) Whole-mount immunostaining of 27 hpf wt and *col15a1b*$^{-/-}$ embryos with guinea pig antibodies directed against rNC1 (left) and with rabbit anti-NC10 (Guillon et al, 2016) as positive control (right).

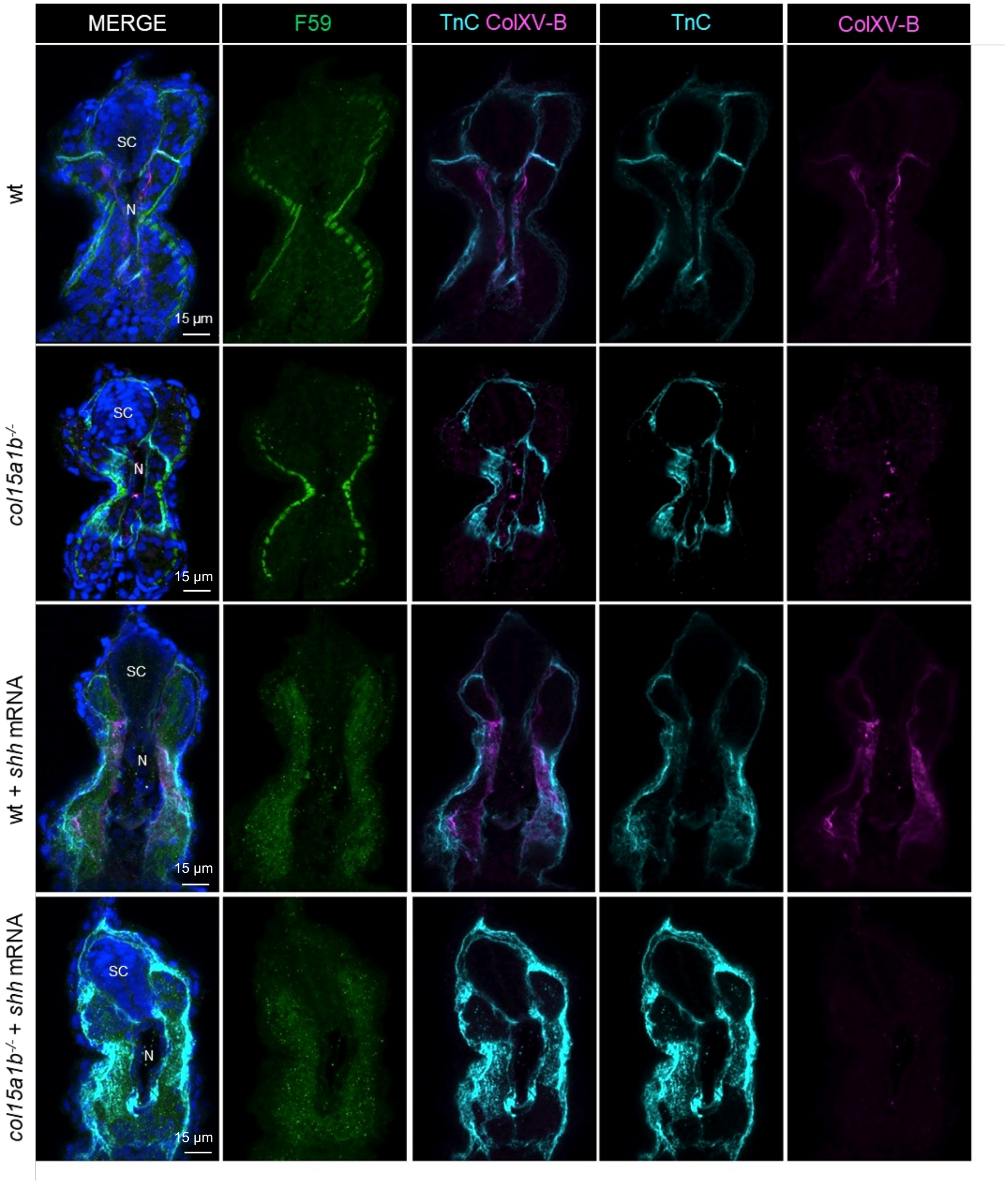


**Figure S3.** Cryosections of 27 hpf wt, *col15a1b*$^{-/-}$, wt *shh*-injected and *col15a1b*$^{-/-}$ *shh*-injected embryos after whole-mount immunostaining with F59 monoclonal antibody (slow muscle marker, green), rabbit anti-TnC (cyan) and guinea pig anti-rNC1 (ColXV-B, magenta)). N, notochord; SC, spinal cord.

A

B

C

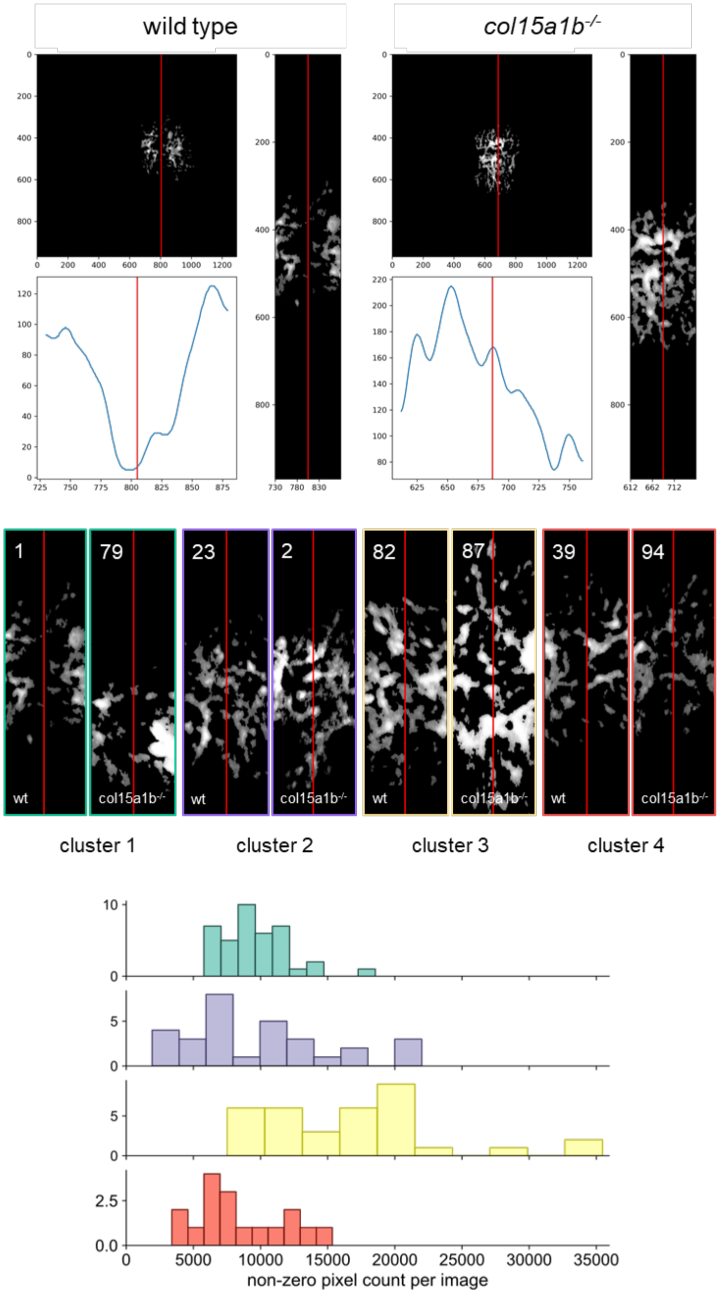


**Figure S4**. (A) Three-plot representation for two embryos: wild type and *col15a1b*$^{-/-}$*.* The top left graph is the 8-bit image resulting from the tenascin-C deposition assessment. The right graph is the 8-bit image resulting from the truncation step of the TnC channel organization analysis. The bottom left graph shows the TnC distribution along the motor path (blue curve) estimated by vertically summing non-zero pixels then smoothed by a Gaussian filter. In all graphs, the red line marks the center of the channel. (B) 8-bit images resulting from the truncation step of the TnC channel organization analysis in wt and *col15a1b*$^{-/-}$ embryos for each cluster. Numbers correspond to the image identifier in the image bank. (C) Histograms of non-zero pixel counts per image per cluster.

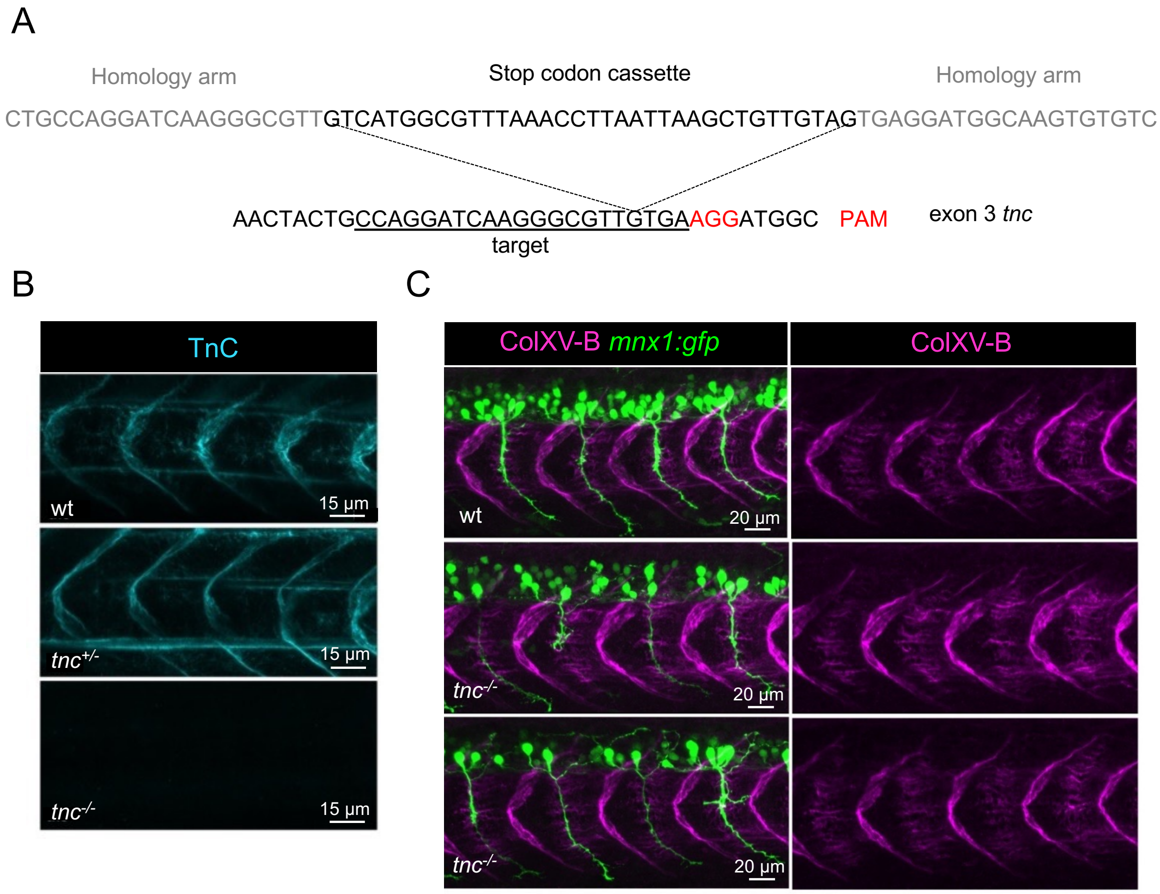


**Figure S5.** Generation of a knock-out line for *tnc* using CRISPR/Cas9 method. (A) Schematic representation of the location of the stop codon cassette insertion in *tnc* exon 3. Cassette insertion is predicted to result in TnC truncation in the first EGF-like domain. (B) Whole-mount immunostaining of 27 hpf wt, *tnc*$^{+/-}$ and *tnc*$^{-/-}$ with anti-TnC antibodies. (D) Whole-mount immunostaining of 27 hpf *mnx1:gfp* and *tnc*$^{-/-}$,*mnx1:gfp* embryos with anti-rNC1 (magenta) and anti-GFP (green).

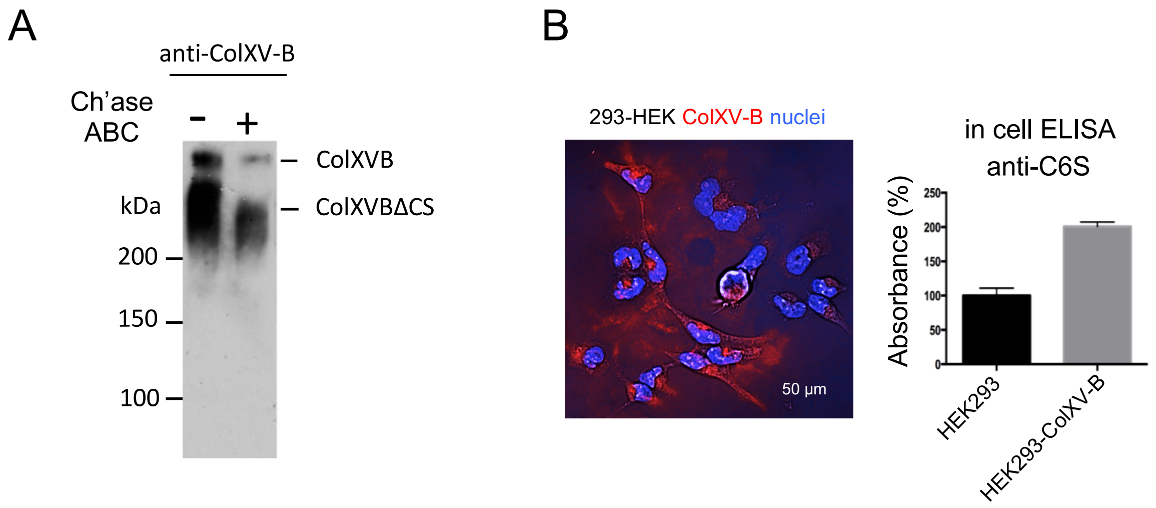


**Figure S6.** ColXV-B carries chondroitin sulfate chains. (A) Western blot analysis with anti-ColXV-B antibody of recombinant ColXV-B incubated with (+) or without (-) chondroitinase ABC (Ch'ase ABC). (B) Immunostaining of ColXV-B transfected HEK293 cells with anti NC10 antibodies (left panel), quantification of chondroitin sulfate chains by absorbance with anti-C6S antibody using in cell ELISA assay at 365 nm. Error bars represent means±SEM (right panel).

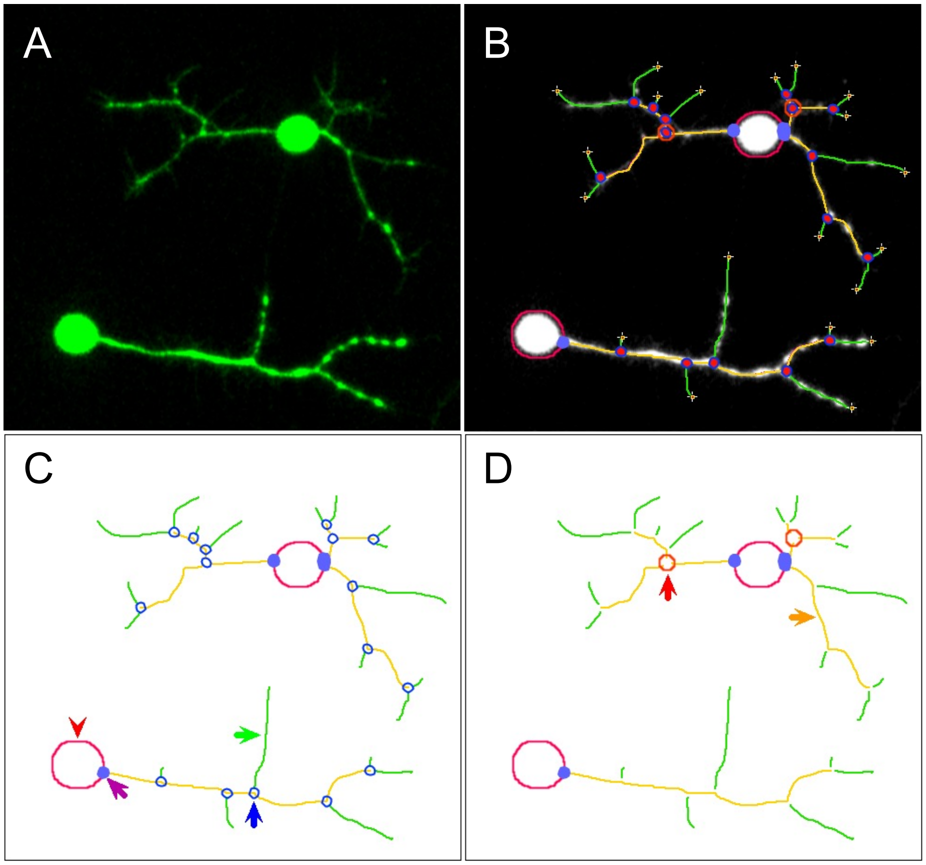


**Figure S7.** In vitro neuronal structure analysis. (A) Initial image sample. (B) Network analysis with customized version of « Angiogenesis Analyzer ». (C) Quantified vectorial objects: red arrow head, « soma »; violet arrow, « anchorage junction »; green arrow, « branch »; blue arrows, « junctions ». (D) Red arrow, « master junction »; orange arrow, « master segment ».

**Supplementary Tables**

| SLOW MUSCLE PRECURSORS – CLUSTER 10 | | | | | |
|---|---|---|---|---|---|
| CORE MATRISOME | | | | | |
| Collagens | | ECM glycoproteins | | Proteoglycans | |
| *col4a1* | 0.33 | *agrn* | 0.56 | | |
| *col4a2* | 0.27 | *efemp2a* | 0.83 | | |
| *col5a1* | 0.7 | *efemp2b* | 1.82 | | |
| *col5a2a* | 0.4 | *fn1a* | 0.37 | | |
| | | *fn1b* | 1.8 | | |
| | | *igfbp5b* | 0.46 | | |
| | | *mfap1* | 0.54 | | |
| | | *mfap2* | 1.18 | | |
| | | *mfge8b* | 0.27 | | |
| | | *nid2a* | 1.07 | | |
| | | *ntn1a* | 0.27 | | |
| | | *sparc* | 0.27 | | |
| MATRISOME-ASSOCIATED | | | | | |
| Secreted Factors | | ECM Regulators | | ECM-affiliated Proteins | |
| *angptl4* | 0.6 | *ctsa* | 0.3 | *anxa11a* | 0.49 |
| *angptl7* | 0.72 | *ctsba* | 0.3 | *lgals3a* | 0.56 |
| *bmp4* | 0.26 | *ctsd* | 0.69 | *lgals3b* | 0.73 |
| *btc* | 0.32 | *ctsla* | 1.3 | *sdc2* | 0.27 |
| *cxcl12a* | 0.66 | *hpse* | 0.25 | *sdc4* | 0.96 |
| *cxcl12b* | 1.28 | *mmp2* | 0.28 | | |
| *egfl6* | 0.5 | *mmp14a* | 0.54 | | |
| *fgf8a* | 0.37 | *p4ha1a* | 0.28 | | |
| *fgf17* | 0.43 | *p4ha2* | 0.43 | | |
| *fstl1a* | 1.1 | *plod3* | 0.38 | | |
| *fstl1b* | 1.2 | *serpinh1a* | 0.44 | | |
| *hcfc1a* | 0.47 | *serpinh1b* | 3.37 | | |
| *hcfc1b* | 0.27 | *sulf1* | 0.5 | | |
| *ism1* | 0.53 | | | | |
| *mdka* | 2.11 | | | | |
| *ngfb* | 0.48 | | | | |
| *wif1* | 0.37 | | | | |

**Table S1.** Classification into matrisome categories of the matrisome genes expressed by cluster 10 SMPs and their average expression (log-normalized expression values).

| SLOW MUSCLE PRECURSORS – CLUSTER 11 | | | | | |
|---|---|---|---|---|---|
| CORE MATRISOME | | | | | |
| Collagens | | ECM glycoproteins | | Proteoglycans | |
| *col4a1* | 0.42 | *agrn* | 0.34 | *hapln1a* | 1.17 |
| *col4a2* | 0.34 | *efemp2a* | 2.96 | *vcana* | 0.66 |
| *col5a1* | 1.16 | *efemp2b* | 1.67 | | |
| *col5a2a* | 0.57 | *fbn2b* | 0.49 | | |
| *col15a1b* | 0.45 | *fn1b* | 1.12 | | |
| *col18a1a* | 0.57 | *fras1* | 0.35 | | |
| | | *igfbp5b* | 0.61 | | |
| | | *lamc1* | 0.37 | | |
| | | *mfap1* | 0.45 | | |
| | | *mfap2* | 1.01 | | |
| | | *mfge8b* | 0.67 | | |
| | | *nid2a* | 1.38 | | |
| | | *ntn1a* | 1.13 | | |
| | | *ntn5* | 0.41 | | |
| | | *spon1b* | 0.35 | | |
| | | *tnc* | 0.46 | | |
| MATRISOME-ASSOCIATED | | | | | |
| Secreted Factors | | ECM Regulators | | ECM-affiliated Proteins | |
| *angptl4* | 0.6 | *ctsd* | 0.59 | *anxa11a* | 0.75 |
| *btc* | 0.71 | *ctsla* | 0.92 | *anxa11b* | 0.28 |
| *cxcl12a* | 1.28 | *mmp14a* | 0.29 | *anxa13* | 0.46 |
| *cxcl12b* | 1.22 | *p4ha1a* | 0.35 | *frem3* | 0.45 |
| *egfl6* | 0.74 | *p4ha2* | 0.7 | *lgals3a* | 0.68 |
| *fgf17* | 0.56 | *plod3* | 0.44 | *lgals3b* | 0.84 |
| *fstl1a* | 2.23 | *serpinh1a* | 0.38 | *lman1* | 0.36 |
| *fstl1b* | 2.01 | *serpinh1b* | 3.35 | *plxna3* | 0.28 |
| *hbegfa* | 0.5 | *sulf1* | 0.67 | *plxnb1a* | 0.33 |
| *hhip* | 0.53 | | | *sdc4* | 1.09 |
| *inhbb* | 0.28 | | | | |
| *ism1* | 0.5 | | | | |
| *mdka* | 1.16 | | | | |
| *ngfb* | 0.5 | | | | |
| *tnfsf10l* | 0.28 | | | | |
| *wif1* | 0.64 | | | | |

**Table S2.** Classification into matrisome categories of the matrisome genes expressed by cluster 11 SMP and their average expression (log-normalized expression values).

| SLOW MUSCLE PRECURSORS – CLUSTER 5 | | | | | |
|---|---|---|---|---|---|
| CORE MATRISOME | | | | | |
| Collagens | | ECM glycoproteins | | Proteoglycans | |
| *col4a1* | 0.8 | *cthrc1a* | 0.4 | *hapln1a* | 0.9 |
| *col4a2* | 0.67 | *efemp2a* | 2.9 | *optc* | 0.29 |
| *col5a1* | 0.99 | *efemp2b* | 1.4 | *vcana* | 0.4 |
| *col5a2a* | 0.66 | *fbn2b* | 0.29 | | |
| *col11a1a* | 0.4 | *fn1b* | 0.5 | | |
| *col15a1b* | 0.68 | *fras1* | 0.45 | | |
| *col18a1a* | 0.55 | *igfbp1a* | 0.62 | | |
| *col19a1* | 0.4 | *igfbp5b* | 0.48 | | |
| | | *lama5* | 0.26 | | |
| | | *lamc1* | 0.25 | | |
| | | *mfap1* | 0.4 | | |
| | | *mfap2* | 0.7 | | |
| | | *mfge8b* | 0.6 | | |
| | | *mxra5b* | 0.25 | | |
| | | *nid2a* | 0.76 | | |
| | | *ntn1a* | 0.48 | | |
| | | *sparc* | 0.43 | | |
| | | *tnc* | 1.01 | | |
| MATRISOME-ASSOCIATED | | | | | |
| Secreted Factors | | ECM Regulators | | ECM-affiliated Proteins | |
| *btc* | 0.67 | *ctsd* | 0.5 | *anxa11a* | 0.83 |
| *egfl6* | 0.73 | *ctsla* | 0.8 | *anxa13* | 0.64 |
| *fgf17* | 0.29 | *kazald3* | 0.47 | *frem3* | 0.5 |
| *fstl1a* | 1.9 | *p4ha1a* | 0.32 | *gpc4* | 0.37 |
| *fstl1b* | 1.78 | *p4ha2* | 0.5 | *lgals3a* | 0.79 |
| *hbegfa* | 0.74 | *plod3* | 0.5 | *lgals3b* | 0.6 |
| *mdka* | 1.2 | *serpinh1a* | 0.4 | *lman1* | 0.7 |
| *ngfb* | 0.25 | *serpinh1b* | 2.6 | *plxnb1a* | 0.32 |
| *vegfaa* | 0.28 | *sulf1* | 0.64 | *sdc4* | 1.1 |

**Table S3.** Classification into matrisome categories of the matrisome genes expressed by cluster 5 SMP and their average expression (log-normalized expression values).

| SLOW MUSCLE PRECURSORS – CLUSTER 1 | | | | | |
|---|---|---|---|---|---|
| CORE MATRISOME | | | | | |
| Collagens | | ECM glycoproteins | | Proteoglycans | |
| *col4a1* | 0.83 | *cthrc1a* | 1 | *hapln1a* | 0.87 |
| *col4a2* | 0.72 | *efemp2a* | 2.65 | *optc* | 0.4 |
| *col5a1* | 0.96 | *efemp2b* | 0.5 | | |
| *col5a2a* | 0.68 | *emilin1b* | 0.29 | | |
| *col11a1a* | 1.03 | *fras1* | 0.48 | | |
| *col15a1b* | 0.54 | *igfbp1a* | 0.48 | | |
| | | *igfbp5b* | 0.33 | | |
| | | *lama5* | 0.32 | | |
| | | *lamc1* | 0.3 | | |
| | | *mfap1* | 0.3 | | |
| | | *mfap2* | 0.34 | | |
| | | *mfge8b* | 0.3 | | |
| | | *mxra5b* | 0.29 | | |
| | | *nid1a* | 0.56 | | |
| | | *nid2a* | 0.35 | | |
| | | *ntn1a* | 0.28 | | |
| | | *sparc* | 0.63 | | |
| | | *tgfbi* | 0.3 | | |
| | | *tnc* | 1.44 | | |
| MATRISOME-ASSOCIATED | | | | | |
| Secreted Factors | | ECM Regulators | | ECM-affiliated Proteins | |
| *btc* | 0.62 | *ctsd* | 0.45 | *anxa6* | 0.51 |
| *egfl6* | 0.46 | *ctsla* | 0.67 | *anxa11a* | 0.97 |
| *fsta* | 0.28 | *hyal1* | 0.3 | *anxa13* | 0.75 |
| *fstl1a* | 1.29 | *kazald3* | 0.44 | *frem3* | 0.61 |
| *fstl1b* | 1.44 | *p4ha1a* | 0.28 | *gpc4* | 0.46 |
| *hbegfa* | 0.9 | *p4ha2* | 0.59 | *lgals3a* | 0.86 |
| *mdka* | 0.7 | *plod3* | 0.63 | *lgals3b* | 0.43 |
| *pdgfab* | 0.45 | *serpinh1a* | 0.41 | *lman1* | 0.44 |
| *sfrp2* | 0.43 | *serpinh1b* | 2.54 | *plxnb1a* | 0.35 |
| | | *sulf1* | 0.54 | *sdc2* | 0.3 |
| | | | | *sdc4* | 0.89 |
| | | | | *sema4ba* | 0.42 |

**Table S4.** Classification into matrisome categories of the matrisome genes expressed by cluster 1 SMP of the and their average expression (log-normalized expression values).

| MUSCLE PIONEER CELLS | | | | | |
|---|---|---|---|---|---|
| CORE MATRISOME | | | | | |
| Collagens | | ECM glycoproteins | | Proteoglycans | |
| *col1a2* | 0.38 | *colq* | 0.26 | *hapln1a* | 1.03 |
| *col4a1* | 0.77 | *cthrc1a* | 0.43 | *optc* | 0.34 |
| *col4a2* | 0.68 | *efemp2a* | 2.82 | | |
| *col5a1* | 0.81 | *efemp2b* | 0.44 | | |
| *col5a2a* | 0.64 | *emilin1b* | 0.29 | | |
| *col11a1a* | 1.02 | *fras1* | 0.48 | | |
| *col15a1b* | 0.7 | *igfbp1a* | 0.74 | | |
| | | *lama5* | 0.46 | | |
| | | *lamc1* | 0.29 | | |
| | | *lgi1a* | 0.38 | | |
| | | *mfap1* | 0.37 | | |
| | | *mfge8b* | 0.34 | | |
| | | *nid1a* | 0.39 | | |
| | | *nid2a* | 0.4 | | |
| | | *ntn1a* | 0.5 | | |
| | | *sparc* | 0.52 | | |
| | | *tgfbi* | 0.28 | | |
| | | *tnc* | 1.78 | | |
| | | *vwa2* | 0.31 | | |
| MATRISOME-ASSOCIATED | | | | | |
| Secreted Factors | | ECM Regulators | | ECM-affiliated Proteins | |
| *btc* | 0.72 | *ctsd* | 0.38 | *anxa6* | 0.45 |
| *egfl6* | 0.54 | *ctsla* | 0.57 | *anxa11a* | 0.9 |
| *fgf13a* | 0.26 | *kazald3* | 0.41 | *anxa13* | 0.79 |
| *fstl1a* | 1.56 | *p4ha1a* | 0.35 | *frem3* | 0.52 |
| *fstl1b* | 1.62 | *p4ha2* | 0.62 | *gpc4* | 0.64 |
| *hbegfa* | 0.98 | *plod3* | 0.52 | *lgals3a* | 0.92 |
| *ism1* | 0.34 | *serpinh1a* | 0.31 | *lgals3b* | 0.48 |
| *mdka* | 0.43 | *serpinh1b* | 2.2 | *lman1* | 0.53 |
| *pdgfab* | 0.32 | *sulf1* | 0.58 | *plxnb1a* | 0.27 |
| *sfrp2* | 0.86 | | | *sdc4* | 0.87 |
| | | | | *sema4ba* | 0.3 |

**Table S5.** Classification into matrisome categories of the matrisome genes expressed by MP and their average expression (log-normalized expression values).
